# Micro Computed Tomography as an Accessible Imaging Platform for Exploring Organism Development and Human Disease Modeling

**DOI:** 10.1101/391300

**Authors:** Todd A. Schoborg, Samantha L. Smith, Lauren N. Smith, H. Douglas Morris, Nasser M. Rusan

**Author notes:** Corresponding Authors Contact Information: Todd Schoborg, Nasser M. Rusan.

## Abstract

Understanding how events at the molecular and cellular scales contribute to tissue form and function is key to uncovering mechanisms driving animal development, physiology and disease. Elucidating these mechanisms has been enhanced through the study of model organisms and the use of sophisticated genetic, biochemical and imaging tools. Here we present an optimized method for non-invasive imaging of *Drosophila melanogaster* at high resolution using micro computed tomography (μ-CT). Our method allows for rapid processing of intact animals at any developmental stage, provides precise quantitative assessment of tissue size and morphology, and permits analysis of inter-organ relationships. We then use the power of μ-CT imaging to model human diseases through the characterization of microcephaly in the fly. Our work demonstrates that μ-CT is a versatile and accessible tool that complements standard imaging techniques, capable of uncovering novel biological mechanisms that have remained undocumented due to limitations of current methods.

## INTRODUCTION

*Drosophila melanogaster* is a foundational model organism that has contributed to our understanding of fundamental cellular and developmental pathways required to build complex multicellular animals. Furthermore, flies have become an important model for over 230 human diseases including cancer, cardiovascular diseases, neurological and metabolic disorders, and others that span virtually every organ system (Millburn et al., 2016). Flies share many of the same genes implicated in human disease, and genome engineering (e.g., P-element transformation & CRISPR/Cas9) can be used to manipulate the *Drosophila* orthologs of these disease genes or introduce human transgenes containing exact patient mutations to better understand disease etiology (Ugur et al., 2016). The powerful genetic tools available in *Drosophila*, coupled with biochemical, molecular, and modern imaging methods has allowed researchers to uncover molecular mechanisms underlying normal development, and how failure of these mechanisms can lead to disease.

Visualization of subcellular, cellular, and tissue organization in the fly relies heavily on fluorescent-based light microscopy such as confocal, which has elucidated countless biological mechanisms in the fly. Other imaging techniques such electron microscopy and super-resolution microscopy, although more challenging, offer additional advantages in resolution. The common disadvantage to these methods is that they do not provide a holistic, whole organism research approach that preserves organ-organ relationships. This type of whole animal analysis is critical for understanding how organisms normally develop and how inter-organ assembly and communication might influence disease states. Thus, there are significant gaps in our understanding of basic developmental mechanisms in flies and other model organisms that could be filled by other imaging technologies.

We sought to adopt and adapt an imaging modality that permits non-invasive imaging of intact animals and incorporate this type of imaging into our current genetic, cell, and developmental research. Several imaging platforms are available for medical research, including magnetic resonance imaging (MRI), optical coherence tomography (OCT), and ultra-microscopy (Holmes, 2009; Jährling et al., 2010; Morton et al., 1990; Null et al., 2008). Our study focused on micro computed X-ray tomography (μ-CT), which has the greatest potential for high-resolution and nondestructive imaging of tissues in a commercially available and readily accessible platform. μ-CT relies on attenuation of X-rays to generate image contrast in acquired radiographs, followed by 3D reconstruction algorithms to produce tomograms (Morton et al., 1990). Pre-clinical μ-CT has proven useful for diagnostic imaging of mammalian disease and genetic models, including mice and rats (Clark and Badea, 2014). μ-CT has also been used for descriptive and comparative analysis of tissue morphology and gross development studies of insect species such as bees (*Apis & Bombyx* sp.), blow flies (*Calliphora* sp.) and hornets (*Vespa* sp.) (Betz et al., 2007; Metscher, 2009; Smith et al., 2016; Sombke et al., 2015; Wipfler et al., 2016). However, μ-CT is grossly underutilized as only a handful of studies have applied μ-CT to *Drosophila melanogaster* biology (Chen et al., 2017; Fabian et al., 2016; Harrison et al., 2017; Klok et al., 2016; Mattei et al., 2015; Mizutani et al., 2013, 2008), and even fewer studies have combined μ-CT with light microscopy, molecular genetics, and other modern cell and developmental biology approaches.

Here we establish μ-CT as a versatile and effective tool to complement confocal microscopy for studying organ development and tissue morphogenesis. We highlight how μ-CT provides a detailed map of the entire intact animal at micrometer resolution across development at an unprecedented level of detail, allowing for unbiased analysis of mutant animals by investigating all tissues, not simply ones predicted to be defective. To reach this level of imaging detail, we provide an optimized labelling and imaging protocol that allows for nondestructive imaging of larval, pupal and adult tissues in their native context. We then demonstrate its ability to precisely quantitate tissue size and morphology compared to standard techniques. Finally, we use μ-CT to characterize a *Drosophila* model of autosomal recessive primary microcephaly (MCPH) by investigating mutations in *Abnormal spindle (asp)* and *WD Repeat Domain 62 (Wdr62)*. We show that the N-terminal 1/4 of Asp is sufficient for proper brain size and morphology. Interestingly, mutations in this region of the human ortholog *(ASPM)* are known to cause human microcephaly, suggesting a mechanism of brain size specification that is conserved from flies to man, thus opening new directions for ASPM-related microcephaly research and brain development in general. These data showcase μ-CT as an accessible component of the model organism research toolkit that is capable of uncovering novel development and disease mechanisms.

## RESULTS

### An optimized protocol for micro computed tomography (μ-CT) of Drosophila melanogaster

Our μ-CT protocol for *Drosophila* draws from previous methods (Smith et al., 2016; Sombke et al., 2015; Swart et al., 2016) but focuses on rapid processing of samples that preserves larval, pupal and adult tissues in as close to their native state as possible (Methods). μ-CT generates an image through the differential attenuation of X-rays as they pass through a rotating sample towards the detector (Figure S1A-C). Since soft biological tissues generally do not attenuate X-rays well enough to permit detailed visualization at standard energy levels (40-60 keV), elements with high atomic numbers such as iodine-iodide (I2KI) and phosphotungstic acid (PTA) are often used to stain tissue prior to X-ray exposure. We find that the best results are obtained when animals are fixed using Bouin’s Solution, washed in an isotonic buffer and labelled with either iodine or PTA prior to scanning with the animals fully hydrated in water. Iodine is used as a standard stain that rapidly (1-2 days) labels soft tissue in intact adults; PTA requires longer incubation times (3-5 days) and mechanical cuticle disruption but does allow sharper visualization of tissue than iodine.

Scan times per animal range from 20 minutes to overnight, depending on the resolution desired. Images suitable for quantitative analysis at lower resolution can be scanned in 20-30 minutes, whereas higher resolution scans that allow for detailed morphological analysis require 8-16 hrs. Image reconstructions using NRecon (Bruker) are achieved in seconds for low resolution scans, or minutes for high resolution scans. 3D rendering and image analysis (segmentation) is performed using a workstation computer outfitted with a suitable GPU graphics card. Using this protocol, we obtain consistent and robust visualization of soft tissue with minimal reagent cost and hands-on time (~15 minutes total) (Video 1 & 2).

### μ-CT visualization of Drosophila larval stages

A key advantage of our optimized protocol is that it works for any *Drosophila* developmental stage. *Drosophila* are holometabolous insects, possessing defined embryonic, larval, pupal and adult stages, each characterized by dramatic changes in morphology as development progresses. Furthermore, many disease phenotypes manifest during periods of rapid cell division and growth, such as those occurring during the larval instars, or during metamorphosis when complex tissue morphogenesis takes place. Larval stages have been difficult to address from a whole organism perspective using light microscopy. Intact larvae cannot be visualized at high resolution due to light scattering effects of the cuticle, and while dissection techniques such as larval fillet can partially overcome these issues, it is limited to a small subset of tissues and does not preserve all organs in their native environment (Parton et al., 2010).

To demonstrate the capabilities of μ-CT for high resolution whole animal imaging, we imaged intact wildtype (*yw*) third instar larvae (Figure 1). The acquired tomograms produce a fully reconstructed animal with sufficient resolution and contrast to easily visualize every internal organ, including those most commonly studied by confocal microscopy such as the imaginal discs, nervous system, gut, and reproductive organs (Figure 1A, Video 3). Additionally, tissues much more difficult to dissect or relevant to human disease models are readily identifiable such as the trachea, Malpighian tubules, fat bodies, body wall muscles and the heart (Figure 1A, Video 3).

**Figure 1.**
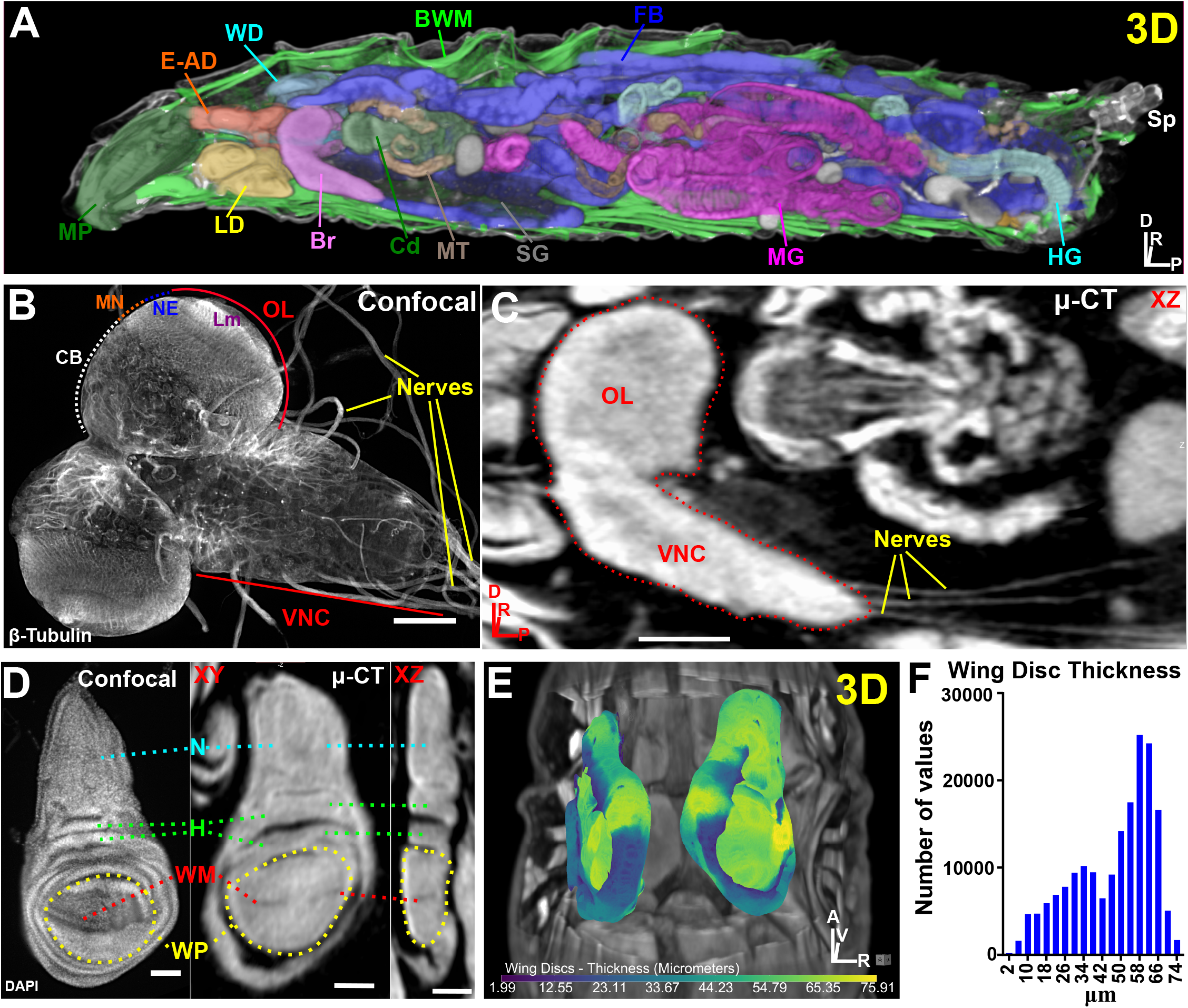
μ-CT reveals all major organs in *Drosophila melanogaster* larvae. (A) 3D display of a larvae optically sliced along the XZ axis showing the MP (Mouthparts); E-AD (Eye-Antennal Discs); WD (Wing Discs); BWM (Body Wall Muscles); FB (Fat Body); Sp (Spiracles); HG (Hindgut); MG (Midgut); SG (Salivary Glands); MT (Malpighian Tubules); Cardia (Cd); Br (Brain); LD (Leg Discs). (B) Confocal image of explanted larval brain labelled with β-tubulin to visualize nerves (yellow lines). Ventral nerve cord (VNC), Optic Lobe (OL), Central Brain (CB), Medulla Neuroblasts (NB), Neuroepithelium (NE) and Lamina (Lm) neuropil are denoted. (C) μ-CT of larval brain viewed along the XZ axis. Note extension of nerves to target organs from the VNC. (D) Confocal image of DAPI-stained wing disc (left) and a μ-CT imaged wing disc (right). Notum (N), hinge (H), wing margin (WM) and wing pouch (WP) are labelled; also note epithelial folds in XZ. (E) 3D representation of wing disc position within the body cavity. Discs are colored by thickness (μm) (see Methods). (F) Histogram of wing disc thickness values. Axis denotes origin, Anterior (A); Posterior (P); Dorsal (D); Ventral (V); Right (R). Scale bars: (B) (C) = 100 μm; (D) = 50 μm. Stain: 0.1N iodine and imaged at 2 μm resolution.

Beyond identification of the major organs, many tissue details clearly emerge. For example, microtubule-rich nerves extending from the larval ventral nerve cord (VNC) can be imaged by confocal with β-tubulin antibodies, but information about these structures within the context of the intact animal is lost upon dissection (Figure 1B). In striking contrast, μ-CT imaging allows one to trace nerves extending from the VNC to their target tissues throughout the body cavity (Figure 1C). Another example of the details evident from μ-CT scans are all major features of wing imaginal discs, including the epithelial folds, wing pouch, notum, hinge, and wing margin (Dorsal/Ventral boundary). These features are stunningly similar to that observed by confocal (Figure 1D). A primary advantage of these μ-CT scans is that they provide a qualitative morphological assessment of the wing disc in addition to precise quantitative information related to disc thickness to study forces driving tissue shape (Figure 1E-F). Furthermore, the wing disc’s relationship to other organs (including its attachment to the trachea) and other imaginal discs can be simultaneously assessed to better understand how spatial organization of tissues within the body cavity influences proper patterning and placement of adult structures (Figure 1E).

Collectively, these data demonstrate that a single staining protocol that requires no dissection and minimal effort/bench time provides high quality images to derive quantifiable data and morphological information on all organ systems *in situ*, allowing for detection of defects anywhere in developing larvae.

### μ-CT used to study the most difficult stages of development: pupation

Where μ-CT truly excels is for the observation of metamorphosis during pupation (Figure 2, Video 4-6). The rapid morphological changes during pupal development, coupled with the histolysis of larval tissues, makes dissection of intact organs for detailed analysis extremely difficult, sometimes impossible. As a result, many pupal tissues remain highly understudied and therefore our knowledge of the mechanisms driving their transition to adult structures is far from complete. This presents an exciting opportunity for discovery when considering pupation and the pharate adult stage encompass a 5-day window of time, nearly 50% of the pre-adult lifespan.

**Figure 2.**
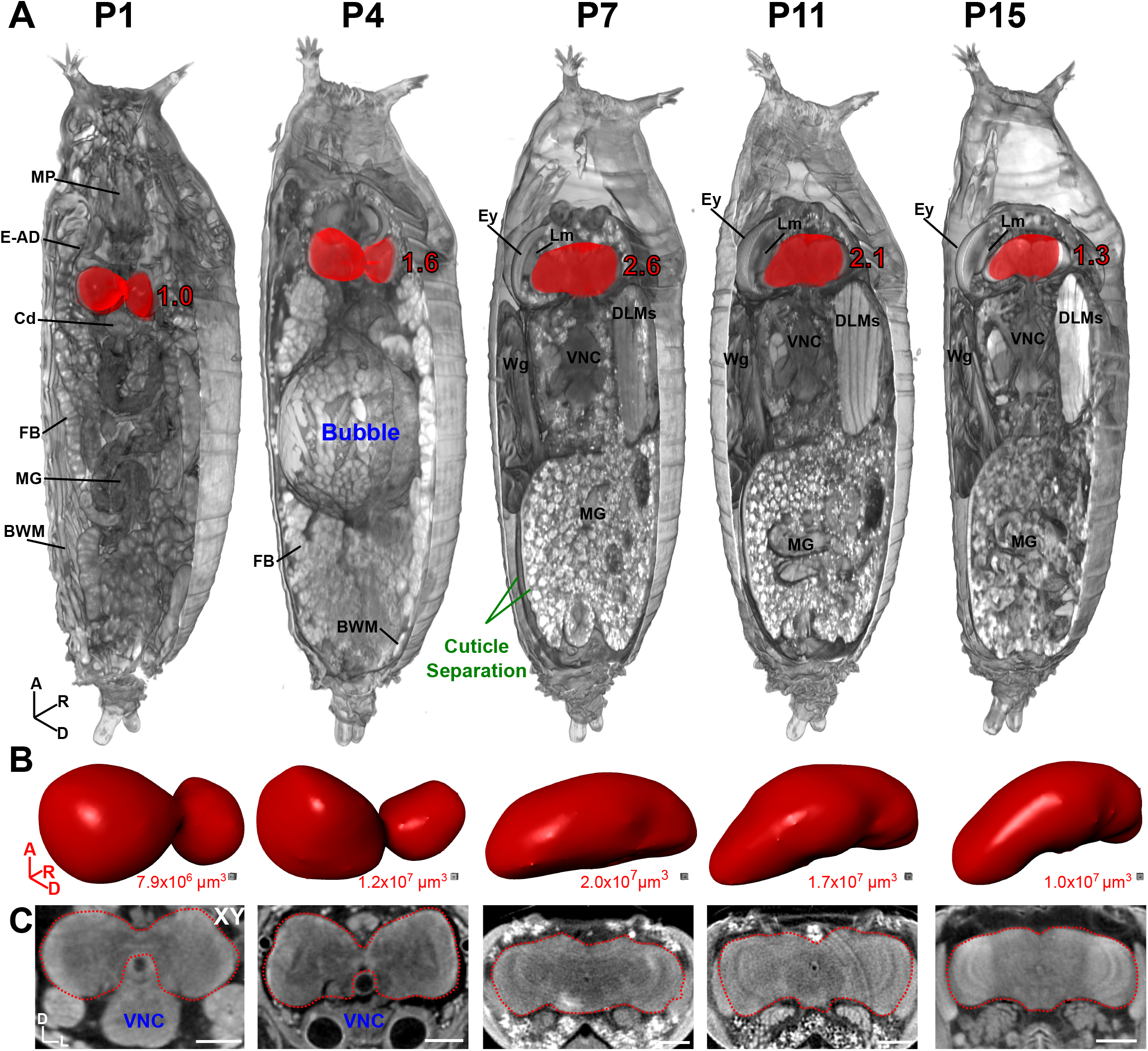
Developmental anatomy of *Drosophila* brains during pupation viewed by μ-CT. (A) Three-dimensional representation of pupae at stage P1, Early P4, P7, P11 and P15. Part of the pupal cuticle and underlying soft tissue were digitally removed to reveal internal structures (axis denotes origin, Anterior (A); Dorsal (D); Right (R)). Brain is highlighted in red; values denote brain size relative to P1. Note the prominent abdominal air bubble at earlyP4 and cuticle separation between the pupal and adult epidermis at the pharate adult stage P7. MP (Mouthparts); E-AD (Eye-Antennal Discs); LD (Leg Discs); Wg (Wing); Cardia (Cd); BWM (Body Wall Muscles); FB (Fat Body); MG (Midgut); DLM (Dorsal Longitudinal Muscles); Ey (Eye); Lm (Lamina), VNC (Ventral Nerve Cord). (B) Isolated brains from (A), shown in same orientation, rendered as a 3D surface. Volume measurements are given below. (C) 2D XY anterior view of brains in (A), axis is denoted D (Dorsal) and L (Left). Scale bars = 100 μm. Stained with 0.1N iodine and imaged at 1.4 μm resolution.

To highlight the capabilities of μ-CT in the study of metamorphosis, we focused on the pupal brain which undergoes dramatic morphological changes between larval and adult stages. Our μ-CT analysis revealed highly dynamic changes in brain morphology and volume throughout this period, increasing 2.6x in size from P1 to P7 (pharate adult) before decreasing to a final volume of 1.3x at P15 prior to eclosion (Figure 2, S2A-B; Video 4 & 5). Furthermore, the distinctive morphological features of pupae described by Bownes and Bainbridge, such as the air bubble at P4 and full separation of the pupal and adult cuticle that marks the pharate adult stage, are clearly evident (Figure 2, S2A-B) (Bainbridge and Bownes, 1981).

In addition to normal development, μ-CT is advantageous for the analysis of mutations that cause pupal lethality (over 2500 alleles listed on FlyBase). μ-CT allows researchers to define the time window of pupal lethality, in addition to the underlying tissue defects that may be responsible. For example, knockdown of *spc105r* (ortholog of the human kinetochore protein *KNL1*) with a Gal4 driver traditionally used to manipulate the larval mushroom body (*ey^OK107^*) leads to massive pupal lethality. Using μ-CT, we discovered that pupae die as pharate adults during a window spanning P11-P15 (Figure 3, Video 6). Moreover, these pupae fail to make proper head structures, instead possessing a remnant structure attached to the thorax that closely resembles the labellum and esophagus, but no antennae, eyes, ocelli or other mouthparts (Figure 3C). Amazingly, a near-completely developed brain of these mutants was found inside the thorax, located ventrally to the dorsal longitudinal muscles (DLMs) and attached to the ventral nerve cord (VNC) at an orthogonal angle (Figure 3C, D, G, H; Video 6). By carefully investigating the μ-CT reconstructions in 2D and 3D, we were able to distinguish brain regions (central brain and optic lobe) and even individual neuropils such as the medulla (Me), lobula (Lo), lobula plate (LoP), and central complex (Cx), although their morphology was clearly defective as compared to P15 *yw* control brains (Figure 3A-D; Video 5). The optic lobes in the *spc105r* RNAi flies are deformed toward the posterior, possibly influenced and shaped by the confines of the surrounding tissues; we could not readily identify lamina, retina, or ocelli structures associated with the ectopically localized brain. Interestingly, our *spc105r* RNAi phenotypes (lethality at pharate adult, remnant head structure consisting of only a labellum), are similar to those reported for loss of the Pax6 orthologs *eyeless* (*ey*) and *twin of eyeless* (*toy*) in eye-antennal discs (Wang and Sun, 2012; Zhu et al., 2017). Whether the Pax6 animals also possessed an ectopically developing brain in the thorax was not reported, which may have simply been overlooked due to technical limitations.

**Figure 3.**
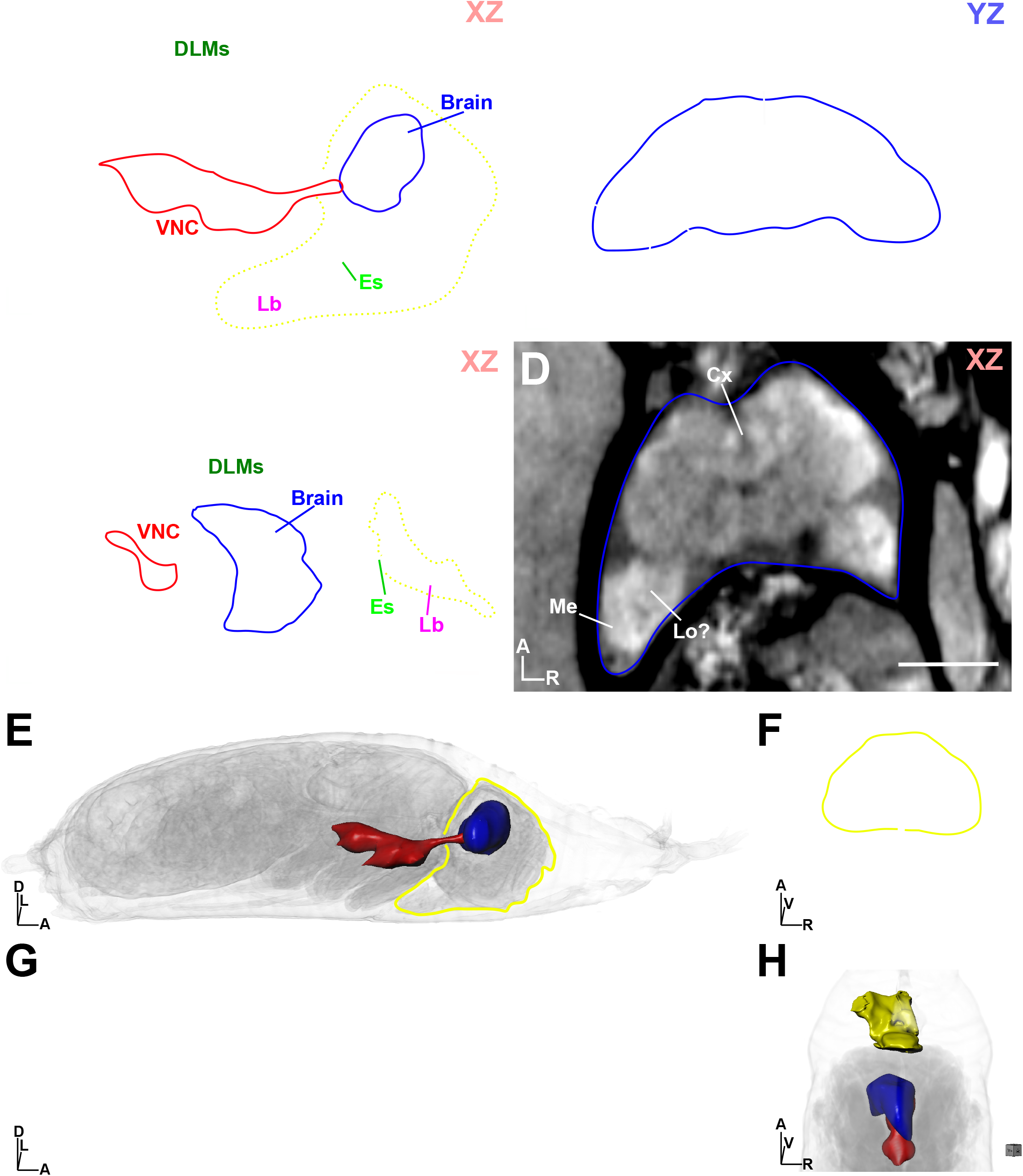
Dissection of a ‘pupal lethal’ mutation by μ-CT. (A) Wildtype (*yw*) pupae at stage P15 shown in XZ orientation. The headcase is outlined in yellow dots, with the ventral nerve cord (VNC) and brain outlined in red and blue, respectively. DLM (Dorsal Longitudinal Muscles); Es (Esophagus); Lb (Labellum). (B) View of the *yw* headcase from (A), viewing the YZ axis from the dorsal perspective. Brain is outlined in blue. Individual brain neuropils are denoted. Me (medulla); Lo (lobula); LoP (lobula plate); Cx (central complex). (C) Analysis of a pupal lethal ‘headless’ fly, resulting from RNAi depletion of *spc105r* (*KNL1*) using the *ey^OK107^* Gal4 driver. View shown is XZ, with the ‘head remnant’ outlined by yellow dots. Note brain (blue outline) location in the thorax; VNC is outlined in red. (D) View of the mutant brain shown in (C); neuropils are highlighted as described in (B). (E) 3D view of the *yw* pupae shown in (A), the headcase is outlined in yellow and the brain (blue) and VNC (red) are rendered as 3D surfaces. (F) Alternative view of the *yw* pupae from (E), viewed from the dorsal perspective. (G) 3D view of the *spc105r* as denoted in (E); head remnant containing the labellum is shown as a yellow surface. (H) Alternative view of the *spc105r* pupae from (G), viewed from the dorsal perspective. Body axes are indicated (D (Dorsal), V (Ventral), A (Anterior), R (Right), L (Left). Scale bars = 100 μm. Stained with 0.1N iodine and imaged at 1.4 μm resolution.

### μ-CT excels as a tool for both morphological and quantitative analysis of Drosophila tissue for modeling human disease

Another advantage of μ-CT is the isotropic pixel resolution of the tomograms, which results in accurate representation of tissue in 3D space and highly precise quantitative measurements. When combined with the morphological information derived from a high resolution μ-CT image, nearly all tissues can be examined for defects in growth, morphogenesis and patterning, not only in the context of normal development and anatomy, but also for the evaluation of relevant human disease phenotypes. We present a few examples of adult tissue analysis:

#### Muscle

Flies serve as an excellent model for identifying muscle-related defects, similar to those found in human patients with muscular dystrophy or other muscle wasting disorders (Kreipke et al., 2017). Fly muscle tissue provides remarkable contrast for μ-CT using either PTA or iodine. Both muscle morphology and quantitative assessment of either 3D muscle volume or 2D crosssectional area of the dorsal longitudinal muscles (DLMs, Figure 4A-B, Video 7) can be used to accurately assess phenotypes in a single scan, as opposed to traditional resin embedding and cross sectioning of the adult thorax which is labor intensive, prone to artifacts, and provides only 2D information.

**Figure 4.**
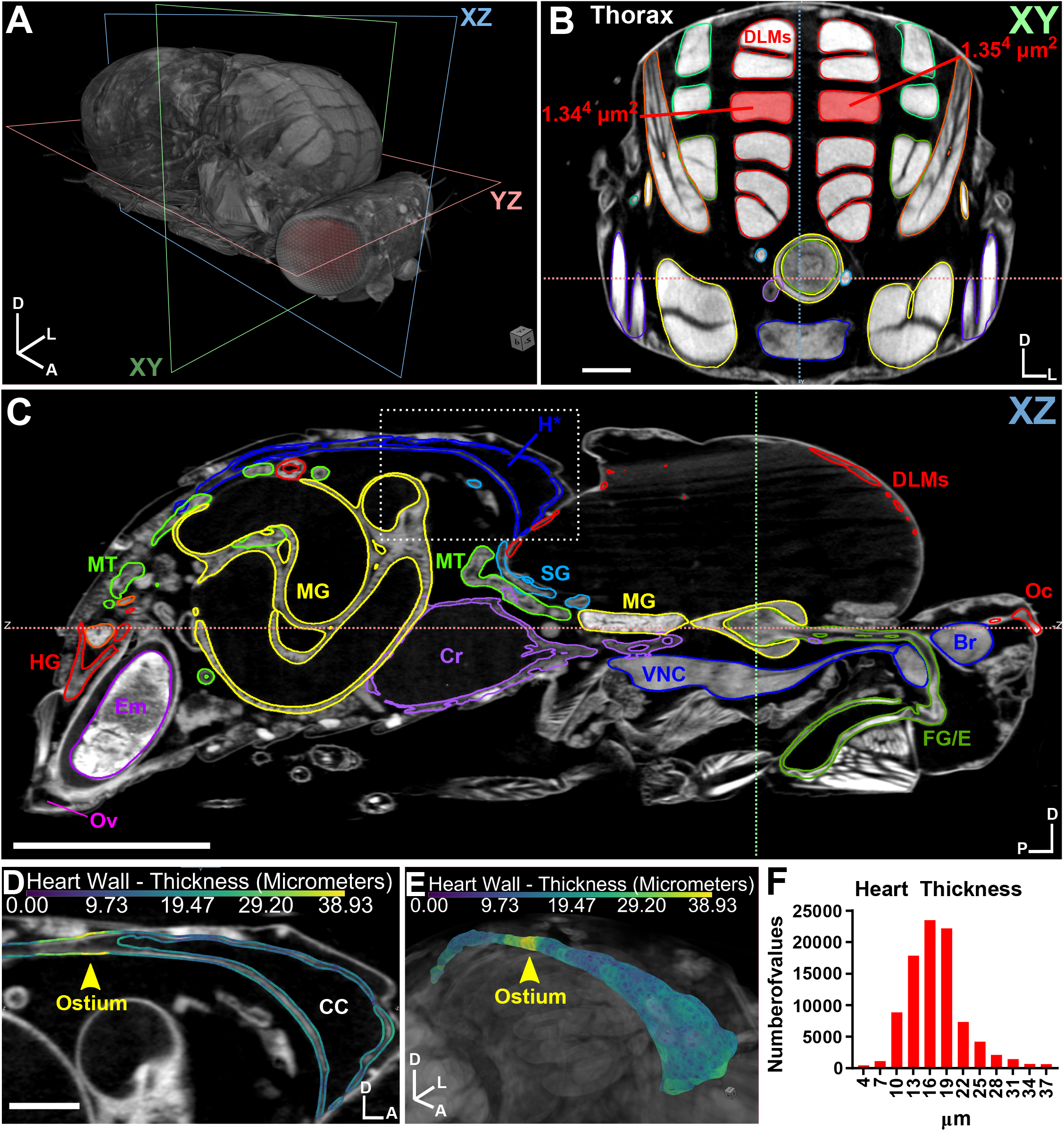
μ-CT of adult *Drosophila melanogaster* highlighting muscle and heart tissue. (A) 3D representation of an adult female fly with the body axis (D (Dorsal), Anterior (A), Left (L)) and imaging planes (XY, XZ, YZ) denoted. (B) 2D XY anterior view of the thorax revealing muscles. A pair of dorsal longitudinal muscles (DLMs) are highlighted and the cross-sectional area is measured. (C) XZ view, organs are segmented by color. HG (Hindgut); MT (Malpighian Tubule); Ov (Ovipositor); Em (Embryo); MG (Midgut); Cr (Crop); H (Heart); SG (Salivary Gland); VNC (Ventral Nerve Cord); FG/E (Foregut/Esophagus); DLMs (Dorsal Longitudinal Muscles); Br (Brain); Oc (Ocelli). (D) Close up of boxed region in (C) revealing the heart. Heart wall thickness (μm) is indicated by color code (see Methods). Position of an ostium is shown (arrowhead); CC (Conical Chamber). (E) 3D representation of the heart, colored by thickness (μm). (F) Histogram of heart thickness values. Body axes are indicated (D (Dorsal), A (Anterior), P (Posterior), L (Left). Scale Bars: (B) (D) =100 μm; (C) = 500 μm. Stained with 0.1N iodine and imaged at 1.25 μm resolution.

#### Heart

The fly heart has also become a reliable system to study human cardiomyopathies. The adult heart tube consists of only two cell types (cardiomyocytes and pericardial cells), while extracellular matrix (ECM), the ventral longitudinal muscles, and alary muscles providing structural support (Rotstein and Paululat, 2016). We used μ-CT to assess heart morphology and thickness to evaluate both the integrity of the heart wall and the ostia, valve-like structures that direct hemolymph flow in the body cavity (Figure 4C-F, Video 8). We found μ-CT to be sensitive enough to easily detect and discriminate between the heart wall (16 μm) and the ostia (35 μm), which appear thicker than the heart wall because of the additional ostial cells that form the valve (Wasserthal, 2007). Thus, μ-CT is an excellent imaging tool to detect small changes in thickness (on the order of a single cell diameter) in extremely delicate tissues such as the heart.

#### Reproductive System

The fly has served as an excellent model for the genetic dissection of sexual reproduction, with almost 2500 genes associated with this term listed on FlyBase. We find that μ-CT can provide useful information for both the male female reproductive system (Figures 5 and 6, Videos 9 & 10) to study fundamental biological mechanisms regarding tissue growth/organization and models of human infertility. For example, high resolution μ-CT scans of an adult female provides a wealth of detail and quantitative information about overall ovary organization and individual egg chambers at each stage of development (germarium-stage 14) (Figure 5A-E, Video 9). We can reliably distinguish nurse cell and their polytene chromosomes, along with clear visualization of the oocyte, similar to confocal microscopy using fluorescent markers (Figure 5C-E). However, imaging the intact animal using μ-CT allows the entire reproductive systems to be studied in its natural spatial arrangement, and its association with other structures more difficult to study by confocal microscopy, such as the ovipositor (Figure 4C, OV). It is noteworthy that the female we present here was withholding eggs, as the embryo seen in the ovipositor has already begun the process of gastrulation, which normally occurs hours after egg laying (Figure 4C, Video 1 & 2).

**Figure 5.**
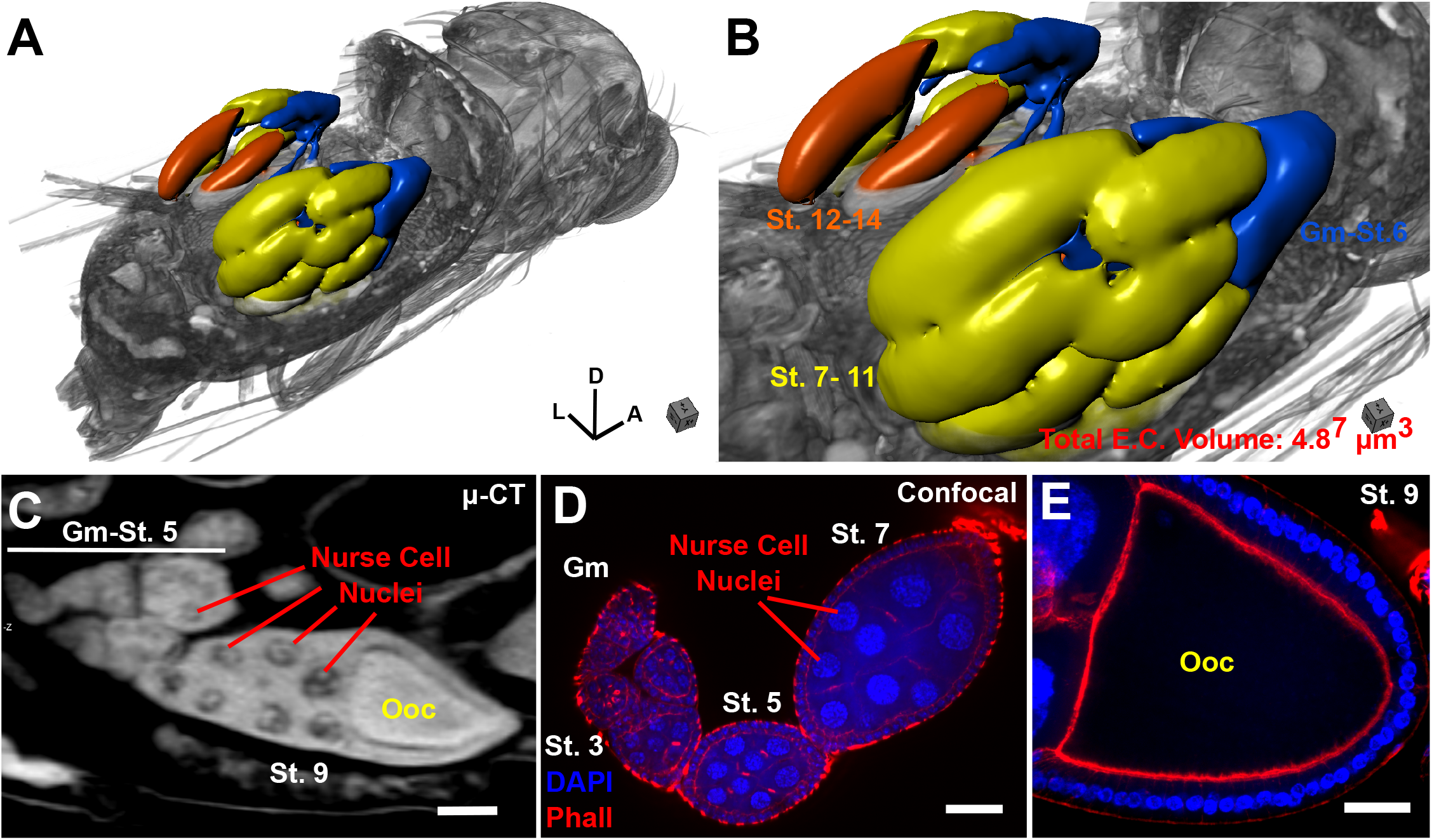
μ-CT of adult *Drosophila melanogaster* highlighting the female reproductive system. (A) 3D view of a female fruit fly, with the abdomen digitally removed to reveal the underlying structure of the ovarioles within the ovary. Body axes are denoted (D (Dorsal), A (Anterior), L (Left). (B) Zoom of (A), ovarioles rendered as a surface and colored according to oogenesis stage (Blue, Germarium (Gm)-Stage 6); Yellow, Stage 7-11; Orange, Stage 12-14). Total egg chamber volume is shown. (C) 2D image of an egg chamber imaged by μ-CT. Stages are denoted (Germarium (Gm)-Stage 5, Stage 9) along with position of the oocyte (Ooc) and nurse cell nuclei (note visualization of polytene chromosomes). (D) Confocal image of an ovariole stained with DAPI and phalloidin, stages are indicated. (E) Confocal image of an oocyte from a stage 9 egg. Scale Bars: (C)-(E) = 50 μm. Stained with 0.1N iodine and imaged at 1.25 μm resolution.

#### Inter-organ relationships

The ability to simultaneously visualize all organs in an intact animal permits the study of interactions between organs, a vastly underappreciated area of research that is critical to understanding the entire phenotypic spectrum of a disease mutation. As an example, left-right asymmetry is a fundamental biological property found throughout metazoans. This is most easily recognized by the human disorder known as situs inversus, in which internal organs are located on the opposite side of the body axis in a mirror-like fashion. In *Drosophila*, this property is exemplified by the directional looping of the spermiduct over the hindgut in a dextral (clockwise) fashion when viewed from the posterior, which occurs during metamorphosis (Adám et al., 2003) (Figure 6, Video 10). μ-CT is perfectly suited to address these types of organ-organ relationships to probe fundamental questions of development, or to ask questions relating to how a diseased tissue can influence the biology of other normal, non-diseased tissues elsewhere in the body.

**Figure 6.**
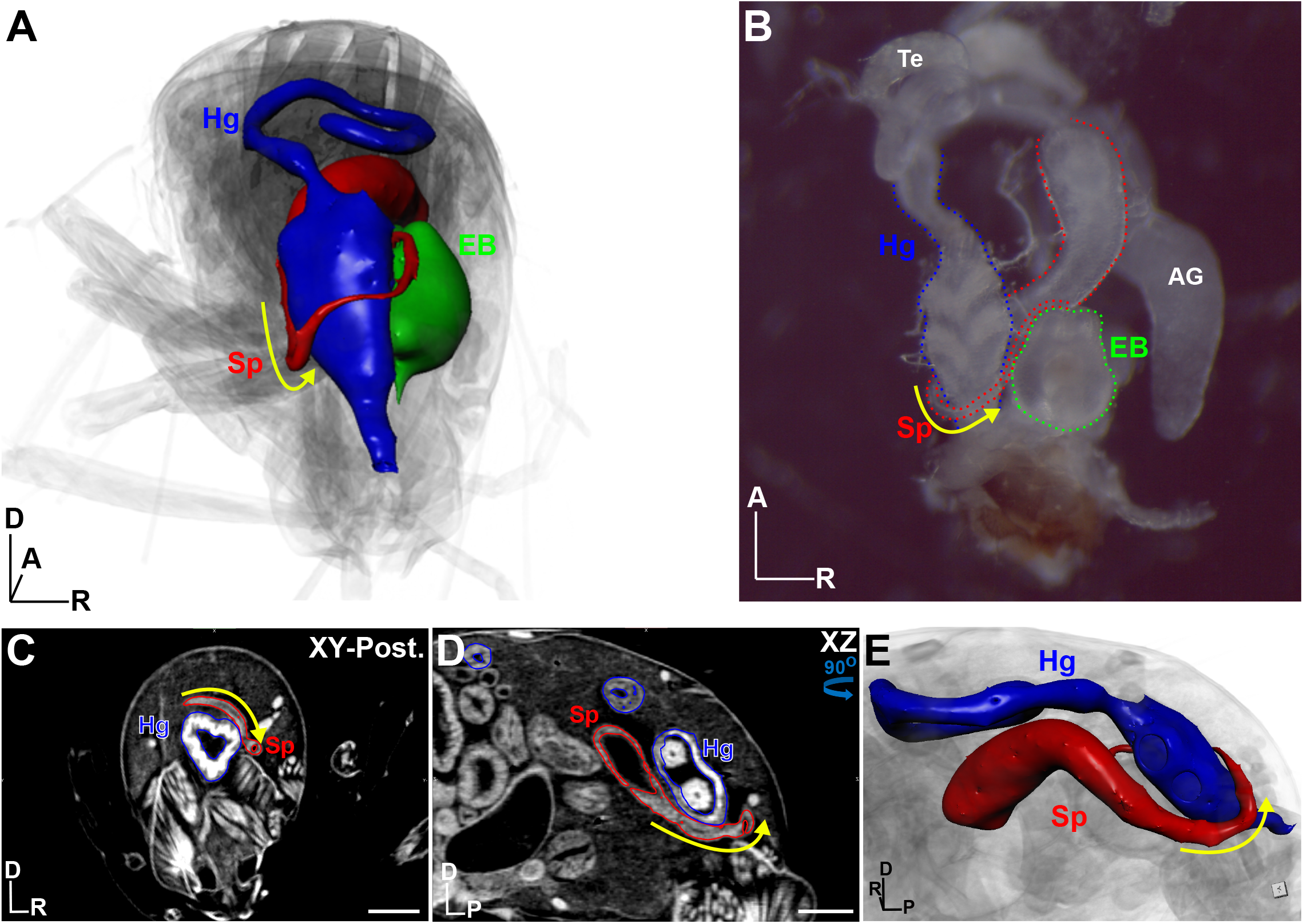
Left-right asymmetry as an example of inter-organ analysis for the study of basic development and human disease. (A) 3D view from the posterior perspective of a male adult fly. The hindgut (Hg, blue), spermiduct (Sp, red) and ejaculatory bulb (EB, green) are rendered as surfaces. Yellow arrow indicates dextral (left-right) looping of the spermiduct over the hindgut into the ejaculatory bulb, located on the right side of the body axis. (B) Dissected version of the organs outlined in (A); testes (te) and accessory glands (AG) are also labelled. (C) XY view from the posterior of the fly shown in (A), the Sp and Hg are outlined. (D) XZ view of the male abdomen shown in (C). (E) 3D view of the Sp and HG as shown in (D), viewed from the left perspective. Yellow arrow denotes dextral looping. Body axes are indicated (A (Anterior), D (Dorsal), P (Posterior), R (Right)). Scale Bars = 100 μm. Flies were stained with 0.1N iodine and imaged at 1.25 μm resolution.

#### Nervous System

As a final demonstration of μ-CT’s ability to derive highly precise quantitative measurements and morphology of tissue, we characterized the adult nervous system in-depth using μ-CT and confocal (Figure 7, Video 11 & 12). Remarkably, most major neuropils designated from confocal imaging of adult brains (Ito et al., 2014) can be assigned in our μ-CT scans of PTA-labelled fly heads (Figure 7A-B, Video 12). However, μ-CT provides additional information about the organization of the visual circuit by allowing for simultaneous observation of the lamina, retina and ocelli. The average entire brain volume of 15 animals measured by μ-CT is 12.9×10^6^ ± 6.0×10^5^ μm^3^ with a coefficient of variation (CV) of 4.7%, compared to confocal measurement of 6.0×10^6^ ± 9.3×10^5^ μm^3^ with a CV of 15.5% (Figure 7C, D). Not only are these μ-CT measurements more precise as shown by lower CV, but they expose an alarming >50% reduction in overall brain volume when measured by confocal imaging. We suspect that dissection and ethanol dehydration required for confocal imaging of chemically cleared brains accounts for most of this volume difference, with a smaller contribution from differences in metal vs fluorescence labelling. To gain additional precision from μ-CT, all measurements can be normalized to thorax width, to account for body size variation (Smith et al., 2016) (Figure 7E & S3; referred to here as the T-ratio). Analysis from the same 15 animals show a T-ratio of 1.6×10^4^ ± 6.4×10^2^ (CV = 4.1%). Thus, we prefer to use T-ratios when performing μ-CT measurements on whole brains and small sub-regions (Figure 7E), as it affords the highest degree of precision.

**Figure 7.**
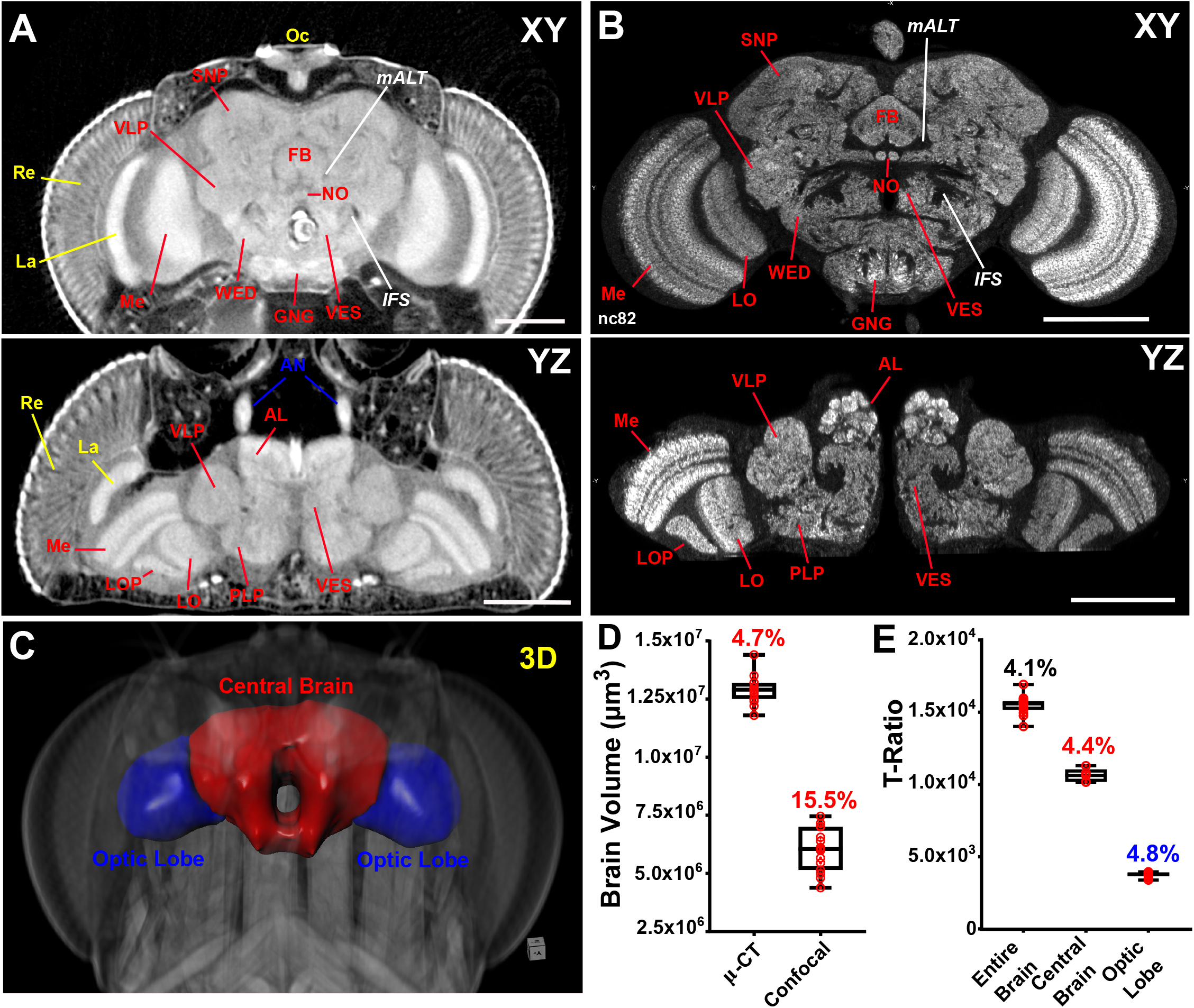
A comparison of μ-CT and confocal microscopy for quantitative and morphological analysis of the nervous system. (A) Anterior XY (top) and Dorsal YZ (bottom) view of an adult brain labeled with PTA and imaged at 700 nm resolution. Locations of individual neuropil structures and fiber tracts are noted. (B) XY (top) and YZ (bottom) view of an adult brain stained for the synapse marker nc82 (α-bruchpilot) and imaged by confocal. Me (Medulla), LO (Lobula), LOP (Lobula Plate), VLP (Ventrolateral Protocerebrum), SNP (Superior Neuropils), WED (Wedge), FB (Fan-shaped body), NO (Noduli), VES (Vest), GNG (Gnathal Ganglia), *mALT* (*medial Antennal Lobe Tract*) and *IFS* (*Inferior Fiber System*), Re (Retina), La (Lamina), Oc (Ocelli). (C) 3D surface representation of the adult brain, indicating the optic lobes (blue) and central brain (red). (D) Comparison of brain volume measurements between μ-CT and confocal microscopy. Box and whisker plots are shown; coefficient of variation (CV%) given. n=15 brains. (E) Measurements of entire brain, central brain and optic lobe volumes normalized to thorax width presented as a T-ratio (Raw brain volume / thorax width). Coefficient of variation (CV%) given. n=5 brains. Scale Bars: 100 μm.

Taken collectively, these data suggest that performing relative measurements of tissues eliminates sampling error, provides an accurate representation of biological variation in tissue size and reduces the required sample size, making μ-CT a preferred tool for quantitative analysis.

### μ-CT for modeling human microcephaly in Drosophila

To demonstrate the utility of μ-CT when combined with the genetic power of *Drosophila*, we applied the technique to better characterize brain defects in two models of human microcephaly, which is characterized by a reduction in brain size, cognitive function and reduced lifespan (O’Neill et al., 2018). We previously showed that mutations in the fly orthologue of *Abnormal Spindle-Like Microcephaly Associated (ASPM)*, known as *Abnormal Spindle (Asp)* recapitulate the microcephaly phenotype seen in human patients, with adult flies displaying a reduction in head and brain size (Schoborg et al., 2015). We used μ-CT to explore heterozygous adult control (*asp^t25^*/+, Figure 8A) and *asp* mutant (*asp^t25^/Df*, Figure 8B) animals. Morphological examination revealed that optic lobe neuropils (medulla, lobula and lobula plate) in *asp* mutants were extremely disorganized while the central brain neuropils appeared only mildly affected compared to wildtype tissue (Figure 8A, B; Figure S4B), a finding supported by confocal imaging (Figure S4A). Unexpectedly, we noted severe defects in the entire visual circuit—the morphology of the lamina and the retina were severely compromised, and the ocelli were either completely absent, extremely disorganized, or reduced in size in *asp* mutant animals. This suggests that *asp* is critical for proper development of the entire visual circuit and the reduction in head size of *asp* mutant animals results from multiple tissue defects, rather than just a small brain *per se* (Figure 8A, B, Video 13 & 14). It also serves as another example of how μ-CT serves as an effective tool for unbiased discovery of novel or overlooked mutant phenotypes, which is critical for understanding disease etiology.

**Figure 8.**
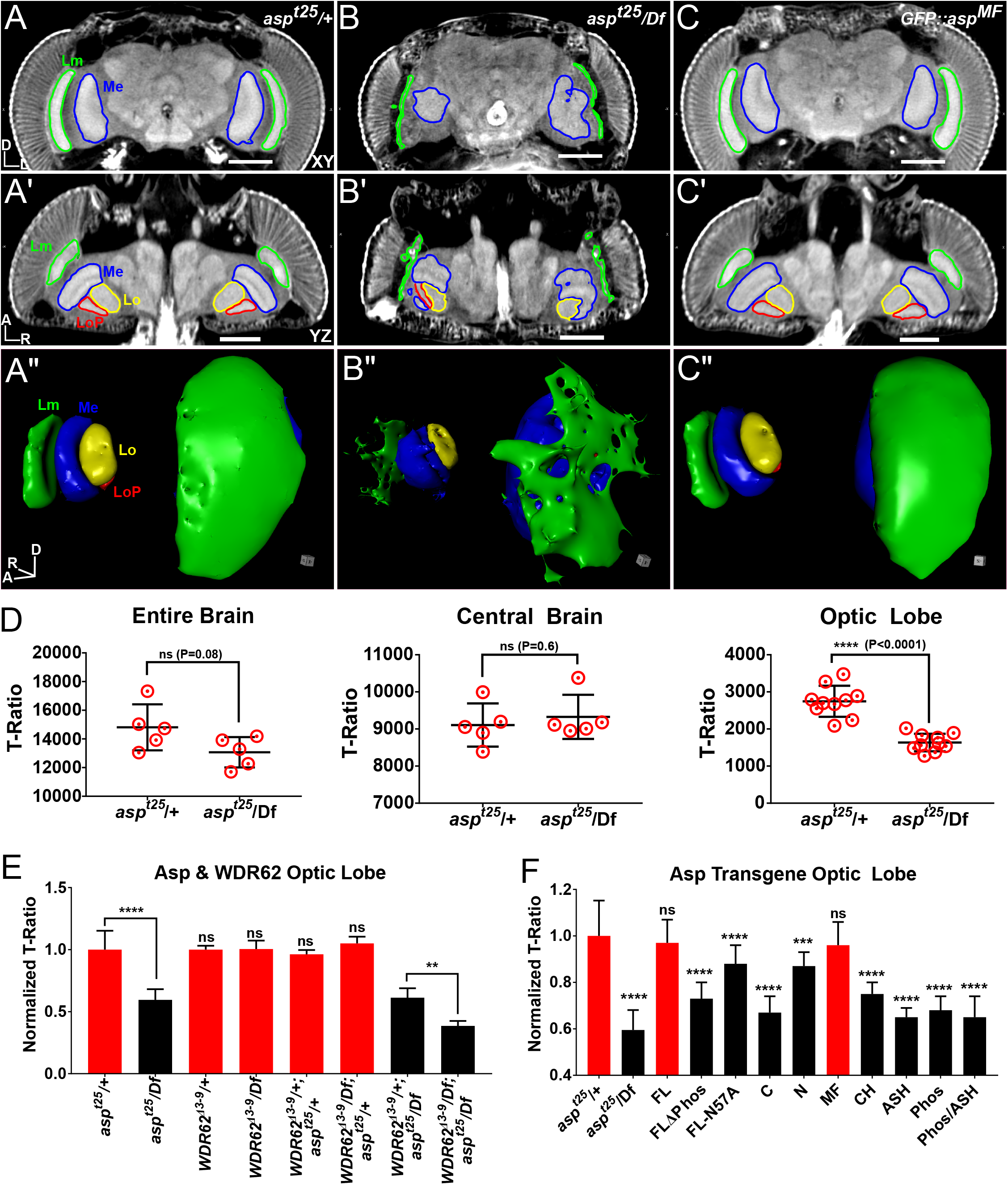
Autosomal Recessive Primary Microcephaly (MCPH) modeled in the adult fly. μ-CT scans of adult heads labelled with PTA and scanned at 1.25 μm resolution from (A, A’, A”) heterozygous control (*asp^t25^*/+), (B, B’,B”) *asp* mutant (*asp^t25^/Df*), and (C,C’,C”) *GFP::asp^MF^* transgenic rescue flies. (A,B,C) Anterior XY view; (A’,B’,C’) Dorsal YZ view; (A”,B”,C”) 3D rendering of visual system. Medulla (Me), lamina (Lm), lobula (Lo) and lobula plate (LOP) neuropils outlined. (D) T-ratio analysis of *asp* brain volume for either the entire brain, central brain only, and optic lobe only. (E) Optic lobe volume of *asp* and *wdr62* brains expressed as a T-Ratio, normalized to *wdr62^Δ3-9^/+; asp^t25^/+* which was set to 1.0. (D) Optic lobe volume of each *GFP::asp* transgene in the *asp^t25^/Df* mutant background. Data is expressed as a T-Ratio, normalized to the *GFP::asp/+; asp^t25^/+* control which was set to 1.0. n= *5* brains, unpaired Student’s t-test. *ns, P*>0.05; ** *P*≥0.01; *** *P*≥0.001; **** *P*≥0.0001. Significance levels shown were derived from direct pairwise comparison between the control (*GFP::asp/+; asp^t25^/+*) and the mutant (*GFP::asp/+; asp^t25^/Df*), as shown in Figures S6 and S7. Body axes are indicated (D (Dorsal), A (Anterior), R (Right), Left (L)). Scale Bars: 100 μm.

We also used μ-CT to acquire precise brain volume measurements of heterozygous adult control (*asp^t25^*/+) and *asp* mutant (*asp^t25^/Df*) animals. 3D tomograms revealed a ~12% reduction in total brain volume in *asp* mutant animals, all of which can be accounted for by a ~30% reduction in optic lobe size (Figure 8D); central brain volume was not affected. Interestingly, animals carrying other *asp* alleles (*asp^E3^ & asp^L1^*) (Rujano et al., 2013) showed nearly an identical volume decrease in the optic lobe, but also a significant decrease in the central brain suggesting allele-specific effects on size (Figure S4C). Alternatively, genetic background differences could be a significant factor, as we have observed when comparing two independent heterozygous control animals, *asp^t25^*/+ & *asp^t25^/TM6B* (Figure S4D).

*Asp* has previously been shown to interact genetically with another microcephaly gene, *Wdr62*, to promote proper brain size in vertebrates (Jayaraman et al., 2016). To test whether this pathway is conserved, we scanned flies carrying a mutation in *WDR62* (Ramdas Nair et al., 2016). Our analysis revealed no detectable reduction in the central brain or the optic lobes in *WDR62^Δ3-9^/Df* animals compared to controls (*WDR62^Δ3-9^/+*) (Figure 8E, Figure S5A-B). These results suggest that the smaller brains reported for larval *WDR62* brains do not lead to microcephaly in adults (Ramdas Nair et al., 2016). However, we did detect a genetic interaction between *WDR62* & *Asp*, as double mutant (*WDR62^Δ3-9^/Df; asp^t25^/Df*) optic lobes showed a further ~25% reduction in volume compared to either the *WDR62^Δ3-9^/+; asp^t25^/Df* or *asp^t25^/Df* mutant alone (Figure 8E). Surprisingly, we also measured a significant decrease in central brain volume for the double mutant animals (Figure S5B), which was not observed in each mutant alone.

Finally, to better understand the regions of Asp required for its ability to promote brain size, we scanned *asp* mutants expressing various N-terminal constructs tagged with GFP (Figure S5C-F). This revealed that the first 573 amino acids (1-573) of Asp were sufficient to rescue the microcephaly phenotype, including both the morphology and size defects (Figure 8A-C, F; Figure S5C-F, Figure S7A, Video 15); we term this fragment Asp^MF^ (Minimal Fragment). We then subdivided this fragment and found that neither Asp^ASH^ (containing an ASH domain), or Asp^Phos^ (containing a predicted CDK1 phosphorylation region (Saunders et al., 1997)), could rescue optic lobe size (Figure 8F). Furthermore, we showed that these domains must be within the same polypeptide, as co-expression of Asp^ASH^ and Asp^Phos^ in the same animal (Asp^ASH^/Asp^Phos^) also failed to rescue. To test necessity of the ASH domain or the phosphorylation region, we generated animals with transgenes expressing full-length Asp carrying either a mutation of a highly conserved asparagine residue present in all ASH domain-containing proteins (Ponting, 2006) (Asp^FL-N57A^), or a deletion of the phosphorylation region (Asp^FLΔPhos^). Both failed to restore optic lobe size to the level of the Asp^FL^ control (Figure 8F, Figure S6B-C), demonstrating that the ASH domain and phosphorylation region are necessary, but not sufficient, for Asp’s function in specifying brain size. Although the precise functions of these two regions remain unknown, it is worth noting that an equivalent deletion of the Asp^MF^ region in the human *ASPM* gene has been identified in an MCPH patient (Nicholas et al., 2009), further highlighting conserved mechanisms of brain development.

## DISCUSSION

This work introduces a powerful imaging technique (μ-CT) to the model organism community that compliments commonly used imaging platforms, such as confocal microscopy, to effectively answer questions regarding anatomy, development and disease that cannot be obtained by current methods. The usefulness of μ-CT for applications in model systems continues to increase as commercial platforms push the resolution limit of the technology (50 nm spatial resolution for the Zeiss XRadia 810 Ultra) and synchrotron facilities devote beamlines specifically for X-Ray imaging. Benchtop scanners such as the one utilized in this study can be obtained for ~$300,000 USD, which is substantially cheaper than high-end confocal microscopes. Other non-invasive imaging methods (MRI, OCT or ultramicroscopy) are either cost-prohibitive, lack commercial potential, require highly specialized expertise or can visualize only one specific tissue. μ-CT will continue to become more accessible as researchers realize the versatility of this technology.

The advantages of μ-CT include: 1) Intact specimens are imaged in a nondestructive manner, allowing the entire animal and all its organ systems to be viewed in their native state. Thus, one powerful aspect of μ-CT is the observation of the interplay *between* different tissues or organ systems, a vastly underappreciated area of study required for a systemic understanding of a genotype or disease mutant. 2) No dissections are required, eliminating tissue damage and allowing very delicate or interconnected tissues to be studied. 3) No tissue-specific reagents, such as antibodies, are required. Even poorly studied systems with few available reagents can be studied at high resolution which can simultaneously reveal defects in other tissues/organs that may have been overlooked in previous studies. 4) Metal labeling is easy to optimize—animals can simply be placed back in stain if sufficient contrast is not achieved after a certain incubation time. 5) Isotropic pixel resolution allows for detection of fine structures in their proper 3D space. Axial resolution (z) is 2-3x worse than in x and y for light microscopy and does not allow for an accurate representation of objects in all dimensions. 6) X-rays are less prone to scattering when passing through biological tissue than light, allowing for visualization of very thick specimens without the need for chemical clearing. 7) Precise quantitative analysis of tissue size through body size normalization, allowing statistical requirements to be met from fewer samples per experiment. This can be extremely advantageous if only a few animals of a given mutant genotype can be recovered.

The primary bottleneck of this technique involves image analysis and segmentation, which can be time consuming to perform manually and requires sufficient computing power and GPU graphics cards for 3D rendering of large datasets. Software such as *Dragonfly* that perform rendering and automate image segmentation via machine learning approaches speed analysis workflows and are preferred for detailed analysis and unbiased identification of phenotypes.

We demonstrated the power of μ-CT by investigating brain development. Other groups have utilized larval brains to investigate microcephaly in the fly using light microscopy (Poulton et al., 2017; Rujano et al., 2013). However, our μ-CT analysis of pupal brain size suggests highly dynamic changes in volume (up to 2.6x) as extensive neurogenesis and neuronal remodeling continues throughout metamorphosis that contributes to final brain size in adults (Homem and Knoblich, 2012). Therefore, the adult is an important endpoint to investigate mechanisms of brain growth control, and μ-CT provides the most precise analysis of tissue size in this regard.

By focusing on adult brains in our investigation of the microcephaly gene *Asp*, we revealed that only ~1/4 of the Asp protein is necessary for proper brain size and morphology, indicating that our knowledge of Asp protein function in cells is in its infancy. Determining the function of the ASH domain, its binding partners and the role of posttranslational modifications in the phosphorylation region will be important future work.

Collectively, our data highlights how μ-CT can be seamlessly integrated with other modern cell and developmental biology methods routinely used by the model research community to uncover novel development and disease mechanisms.

## METHODS

### μ-CT

#### Adult Labeling

5-50 adult flies were anesthetized using CO_2_, transferred to a 1.5ml Eppendorf tube containing 1 ml of 0.5% phosphate buffered saline + Triton-X 100 (0.5% PBST). Tubes were capped, tapped gently on the bench top and incubated for 5 minutes. This step removes wax on the fly cuticle and once complete most of the flies will sink to the bottom of the tube. Flies were then fixed in 1 ml Bouin’s solution (5% acetic acid, 9% formaldehyde, 0.9% picric acid; Sigma Aldrich, St. Lous, MO) for 16-24 hours, washed 3×30 minutes in 1 ml of μ-CT Wash Buffer (0.1M Na2HPO4, 1.8% Sucrose) and stained with 1 ml of a 0.1N solution of iodine-iodide (I2KI, Lugol’s solution) or 0.5% phosphotungstic acid (PTA) solution, diluted in water. I2KI penetrates quicker and can label tissue in 1-2 days without mechanical disruption of adult cuticle, whereas PTA requires 5 or more days plus disruption of the cuticle. This is due to the higher molecular weight and hydrodynamic radius of PTA vs I2KI. However, the differential uptake of PTA by tissue yields better contrast ratios that result in excellent visualization of tissue substructure for finer morphological and anatomical detail. To achieve sufficient brain staining with PTA, the mouthparts were carefully removed with a pair of forceps. Flies were then washed in two changes of ultrapure water and stored at room temperature in ultrapure water until scanned. Samples can be stored indefinitely this way, but we recommend scanning within a month for optimal morphological and quantitative analysis. At no point should flies be put at 4C, as this causes the formation of air bubbles, particularly in the headcase, when flies return to room temperature. The use of ethanol should also be avoided since it causes tissue dehydration, thus altering size and quantitative measurements. It is also recommended that all animal genotypes, including relevant controls, are processed and scanned in the same experiment to ensure the highest confidence when comparing datasets. Finally, this protocol does not require rotation/nutation and can be done simply by incubating flies on the benchtop.

#### Larval/Pupal Labeling

We recommend heating larval and pupal samples in 1 ml of 0.5% PBST to 100C for 20 seconds in a heat block, followed by cooling at room temperature for 5 minutes. This kills animals immediately so that they do not progress to later developmental timepoints. Bouin’s solution can then be added as described for adults, although incubation times should be at least 24 hrs. Animals are then washed 3×30 minutes in μ-CT Wash Buffer (0.1M Na2HPO4, 1.8% Sucrose). Prior to iodine labeling, the larvae and pupae cuticle are poked with a microdissection needle at the anterior and posterior ends, avoiding areas with underlying soft tissue to ensure even tissue labelling. Iodine is then added and the procedure carried out as described for adults.

#### Sample Mounting

There is no defined way to mount samples for scanning and may require resourcefulness on the investigator’s part. The biggest priority is that the sample does not move or dry out during imaging. We use a custom made brass holder with an indention at the tip that perfectly fits a small piece of standard plastic capillary. A P10 pipet tip is then wedged into the capillary tube and filled with water. Parafilm is used to wrap connection points to prevent water leaking into the scanner. Using forceps, flies are then transferred into the pipet tip and a dulled 20-gauge needle is used to gently push the fly downward until it becomes wedged along the wall of the tip and cannot move. A piece of parafilm is used to cover the opening of the pipet tip to prevent evaporation during overnight scans. It is important that the capillary tube & pipet tip are as closely aligned along the long axis of the brass sample holder as possible to avoid wobble during the 360 degree sample rotation, resulting in the highest quality reconstructions (The alignment in Figure S1C is slightly off axis).

#### Scanning

Samples were scanned using a SkyScan 1172 desktop scanner controlled by Skyscan software (Bruker) operating on a Dell workstation computer (Intel Xeon X5690 processor @ 3.47GHz (12 CPUs), 50 GB RAM and an NVIDIA Quadro 5000 (4GB available graphics memory) GPU). X-Ray source voltage & current settings: 40 kilovolts (kV), 110 microamps (μA), 4 watts (W) of power. A Hamamatsu 10Mp camera with a 11.54 μm pixel size coupled to a scintillator was used to collect X-rays converted to photons. Medium camera settings at a resolution of 2.95 μm were used for fast scans (~20 minutes), whereas small camera settings at a resolution of 0.7 μm-2 μm and utilizing 360 degrees of sample rotation were used for overnight scans. Random movement was set to 10 and frame averaging ranged from 5-8. The use of lower accelerating voltages of 40 kV or less reduces image blurring due to electron scattering mechanisms and improves image clarity at higher resolution by reducing the photon energy fraction below peak values, producing an image with contrast that is predominately limited to X-ray absorption.

#### Reconstruction

Tomograms were generated using NRecon software (Bruker MicroCT, v1.7.0.4). Reconstruction is sensitive to the acquired image data including image brightness, contrast and optical bench geometry. Reconstruction parameters may vary by sample preparation and acquisition conditions; therefore, in most cases the process is iterative in setting misalignment compensation values, ring artifact reduction settings and beam hardening compensation values to create the highest quality reconstructions. NRecon allows for individual fine-tuning of each of these components using user-defined parameter steps. For our scans, we utilized a built-in shift correction that uses reference scans to compensate for any sample movement during scanning (i.e., thermal fluctuations or slowly varying sample movements) and any remaining misalignment was manually fine-tuned using the misalignment compensation function(Salmon et al., 2009). Ring artifact correction ranged from 10-20, with beam hardening set to 0%.

### μ-CT Image Analysis & Statistics

Dragonfly (v3.6, Object Research Systems (ORS) Inc, Montreal, Canada, 2018; software available at http://www.theobjects.com/dragonfly) was operated on a Dell workstation (Intel Xeon CPU E5-2623 @ 3.00GHz (16 CPUs), 32 GB RAM, NVIDIA Quadro M2000 GPU (20 GB available graphics memory). Segmentation of structures into individual ROIs was performed manually using thresholded images with the 3D paintbrush function. ROIs were then converted to meshes for quantitative analysis and visualization. The built-in 3D movie maker was used to generate all movies. *Thickness values* for the wing disc and heart (Figures 1E-F; 3D-F) were calculated based on the diameter (μm) of a hypothetical sphere that fits between two boundary points within the tissue, which defines the local thickness at that point (the adult heart wall contains over 90,000 of these points). A LUT colored by thickness (μm) is then used to map the diameter of each sphere within the tissue to provide a visual representation of tissue thickness. The thickness values at each point were also exported as individual values and represented as histograms. FIJI (ImageJ v1.52a, National Institutes of Health, Bethesda, MD) was used for all confocal images (Schindelin et al., 2012). Measurements of brain volume from confocal images was achieved by thresholding brains labelled with DAPI & α-nc82 (to label the outer layer of cell bodies and the internal neuropil) and creating a binary mask of the entire Z-stack, which was then measured using the 3D ImageJ suite(Ollion et al., 2013). *T-Ratio:* We used thorax width to normalize all brain measurements to account for any body size variation between samples.

Thorax width was measured by drawing a line across the widest point of the thorax cuticle when viewed from the YZ imaging plane (Figure S3G). To calculate the T-Ratio, raw brain volume was divided by thorax width for each animal. Normalized T-Ratios were derived by setting the T-Ratio from the relevant control genotype to 1 and normalizing the relevant mutant genotypes accordingly. Statistical analysis and graph generation were performed with GraphPad Prism 7 software (La Jolla, CA). An unpaired, two-tailed Student’s t-test (α=0.05) was used for pair-wise brain size comparisons.

### Immunostaining, Antibodies & Microscopy

Larval wing discs were fixed in 4% PFA for 20 minutes at room temperature, washed in 0.5% PBST and incubated with 1X DAPI for 5 minutes. Larval brains and ovaries were fixed in 9% PFA for 30 minutes at room temperature, washed 3×10min in 0.5% PBST. β-Tubulin antibodies (E7, Developmental Studies Hybridoma Bank (DHSB, Iowa City, IA)) were used at 1:50 overnight at 4C. Phalloidin was used at 1:500 overnight at 4C. Adult brain staining was performed exactly as described by the Janelia FlyLight Project (https://www.janelia.org/project-team/flylight/protocols) and mounted in DPX. α-Bruchpilot (nc82) or α-Cadherin (DN-Ex #8) (DHSB) were used at 1:20 (nc82) or 1:30 (DN-Ex #8). Secondary antibodies conjugated to Alexa Fluor 488, 568 and 647 (Life Technologies) were used at 1:500. Fixed and live confocal imaging of larval wing discs, brains and adult ovaries were performed using a Nikon Eclipse Ti inverted microscope with a 10×/0.30 NA plan Fluor or 40×/1.30 NA plan Fluor objective, a CSU-22 spinning disc module (Yokogawa), an ORCA-Flash4.0 CMOS camera (Hamamatsu) or interline-transfer cooled charge-coupled device camera (CoolSNAP HQ2; Photometrics) and 405 nm, 491 nm, 562 nm and 647 nm solid state illumination lasers (VisiTech International). The microscope was controlled by and images acquired using MetaMorph (v7.7.10, Molecular Devices).

Confocal imaging of adult brains was performed on a Zeiss LSM 880 with Airyscan and imaged using a 63X/1.4NA plan Apo objective, controlled by Zen Black software.

### Asp Transgene Expression Levels

We attempted to measure Asp transgene levels using standard Western Blotting techniques with α-GFP antibodies; however, this approach consistently failed despite numerous attempts to optimize lysis and running conditions(Rujano et al., 2013; Schoborg et al., 2015). Instead we directly measured GFP fluorescence from dissected larval brains that were imaged live to determine expression levels. At least 10 third instar larvae per genotype were washed in a dish of 1X PBS. The brains were dissected in room temperature Schneider’s media with 1X Antibiotic-Antimycotic and kept in media until imaging. Brains were placed into a 25-50 μl droplet of media on the gas-permeable membrane surface of a 50 mm lummox dish (Sarstedt). Four drops of Halocarbon oil 700 (Sigma-Aldrich) were placed on membrane surface to make the corners of a square corresponding to the size of a 22×22-1.5 microscope coverslip (Fisher Scientific). The coverslip was gently placed on top of the sample, and excess media was wicked away using a Kimwipe to flatten and secure the brains. Brains were imaged live on a Nikon Eclipse Ti Spinning Disc confocal with a 10×/ 0.30 NA objective and a 491 nm laser with 2000 ms exposure and 100% laser power. For each data point, a single image was taken of the entire isolated central nervous system (CNS) including the brain and VNC. At least two independent experiments were performed for each genotype. All data analysis was performed using ImageJ. Mean pixel intensity (MPI) was calculated for (1) an entire outline of each *asp* transgenic animal CNS (MPI_CNS_), (2) an area away from the sample for background (MPI_Bkg_), and 3) average MPI for wiltype (*yw*) CNS (MPI_yw_). Final fluorescence measurement MPI_final_ was calculated by MPI_CNS_ - MPI_Bkg_-MPI_yw_. All data was normalized to Asp full-length GFP.

### Fly stocks and husbandry

All stocks and crosses were maintained on standard cornmeal-agar media at room temperature (20-22°C). Fly stocks used were as follows: control (*yw*). *asp^t25^/TM6B, Hu, Tb* (*Schoborg et al., 2015*). *w^1118^; Df*(*3R*)*BSC519/TM6C, Sb*^1^ *cu*^1^ (*asp* deficiency, Bloomington Stock #25023). *w; e11, asp^E3^/TM3, Ser, Act-GFP & asp^L1^/TM6B, Hu, Tb*(*Gonzalez et al., 1998*). *wdr62^Δ3-9^/CyO, GFP* (Cabernard Lab (Ramdas Nair et al., 2016)). *w^1118^; Df*(*2L*)*Exel8005/CyO* (*wdr62* deficiency, Bloomington Stock #7779). *OK107-GAL4* (*w*\*; *P*{*w[+mW.hs]*=*GawB*}*OK107 ey^OK107/In^*(*4*)*ci^D^, ci^D^ pan^ciD^ sv^spa-pol^*; Bloomington Stock #854). *Spc105r RNAi* (*y1 sc* v*^1^; *P*{*TRiP.HMS01752*}*attP2/TM3, Sb^1^*; Bloomington Stock #38534). All *asp* transgenic animals were produced in *yw* genetic background (BestGene Inc, Chino Hills, CA). *asp* transgene genotype combinations were used as follows: Control (*yw*; *GFP-asp*\*/+; *asp^t25^*/+); Mutants (*yw*; *GFP-asp*\*/+; *asp^t25^/aspDf*).

### Pupal Staging

Pupae (*yw*) were staged according to Bainbridge and Bownes (Bainbridge and Bownes, 1981), whose morphological descriptions were accomplished with a “dissecting microscope with lateral illumination close to the stage, interposing the tip of a pair of watch maker’s forceps between the lamp and the puparium to cast a shadow over any feature which may be obscured by surface reflexion but which would show up with light scattered inside the animal”. Remarkably, nearly all these features described over 30 years ago were clearly evident our μ-CT scans, allowing for unambiguous identification of pupal stage.

## ACKNOWLEDGMENTS

We thank Carey Fagerstrom for molecular cloning of Asp transgenes, Brian Galletta for critical reading of the manuscript and help with *asp* larval brain dissections and fluorescent measurements, Ryan O’Neil for critical reading of the manuscript and providing the *spc105^RNAi^* animals, Rachel Ng for assisting with μ-CT scans, Danielle Donahue and Brenda Klaunberg of the NIH Mouse Imaging Facility for μ-CT training, advice and discussion, Mike Marsh from Object Research Systems for *Dragonfly* technical support. This work is supported by the Division of Intramural Research at the National Institutes of Health/National Heart, Lung, and Blood Institute (1ZIAHL006126 to NMR and 1K22HL137902 to TAS).

## AUTHOR CONTRIBUTIONS

TAS devised the project, optimized the μ-CT protocol, performed data collection and image analysis, and wrote the manuscript. SLS and LNS assisted with data collection. HDM performed the initial μ-CT scans and optimized the protocol. NMR supervised the project and wrote the manuscript.

## MATERIALS & CORRESPONDENCE

Requests should be made to either TAS or NMR.

## SUPPLEMENTARY INFORMATION

**Supplementary Figure 1.**
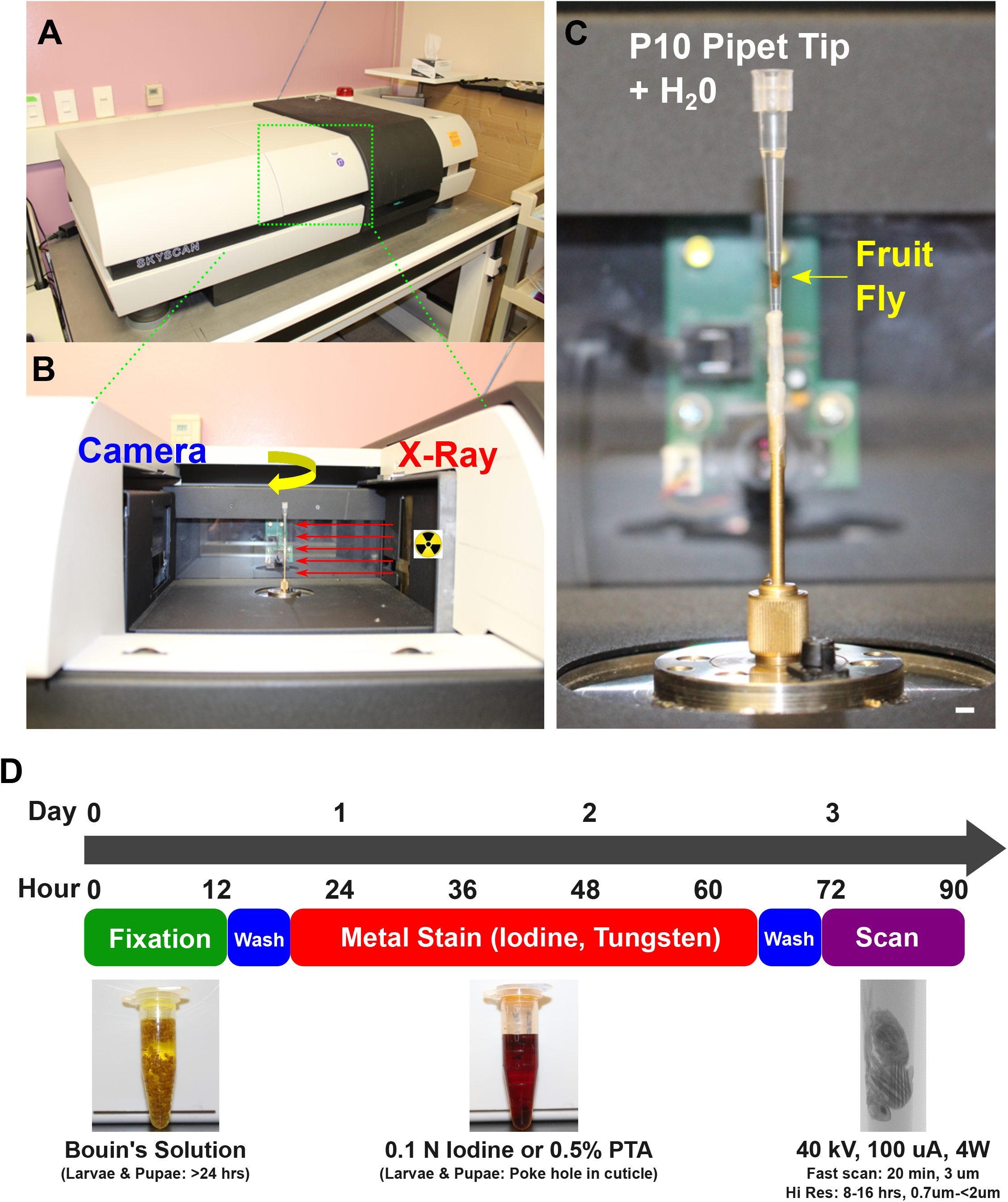
Overview of μ-CT. (A) The benchtop scanner utilized for this study, a Skyscan 1172 (Bruker). (B) Open door view of the imaging path. X-rays are generated by the source at the right and travel towards the camera (left), which detects attenuation of the X-rays as they pass through a rotating sample in order to generate contrast. (C) Close up view of a sample ready for scanning in the rotating sample holder. Position of the fruit fly is indicated by arrow. Note the pipet tip is not perfectly parallel to the long axis of rotation here, which will result in a lower quality reconstruction. (D) Graphical representation of the labelling protocol outlined in *Methods*. Scale Bars (C) = 2 mm.

**Supplementary Figure 2. Related to Figure 2.**
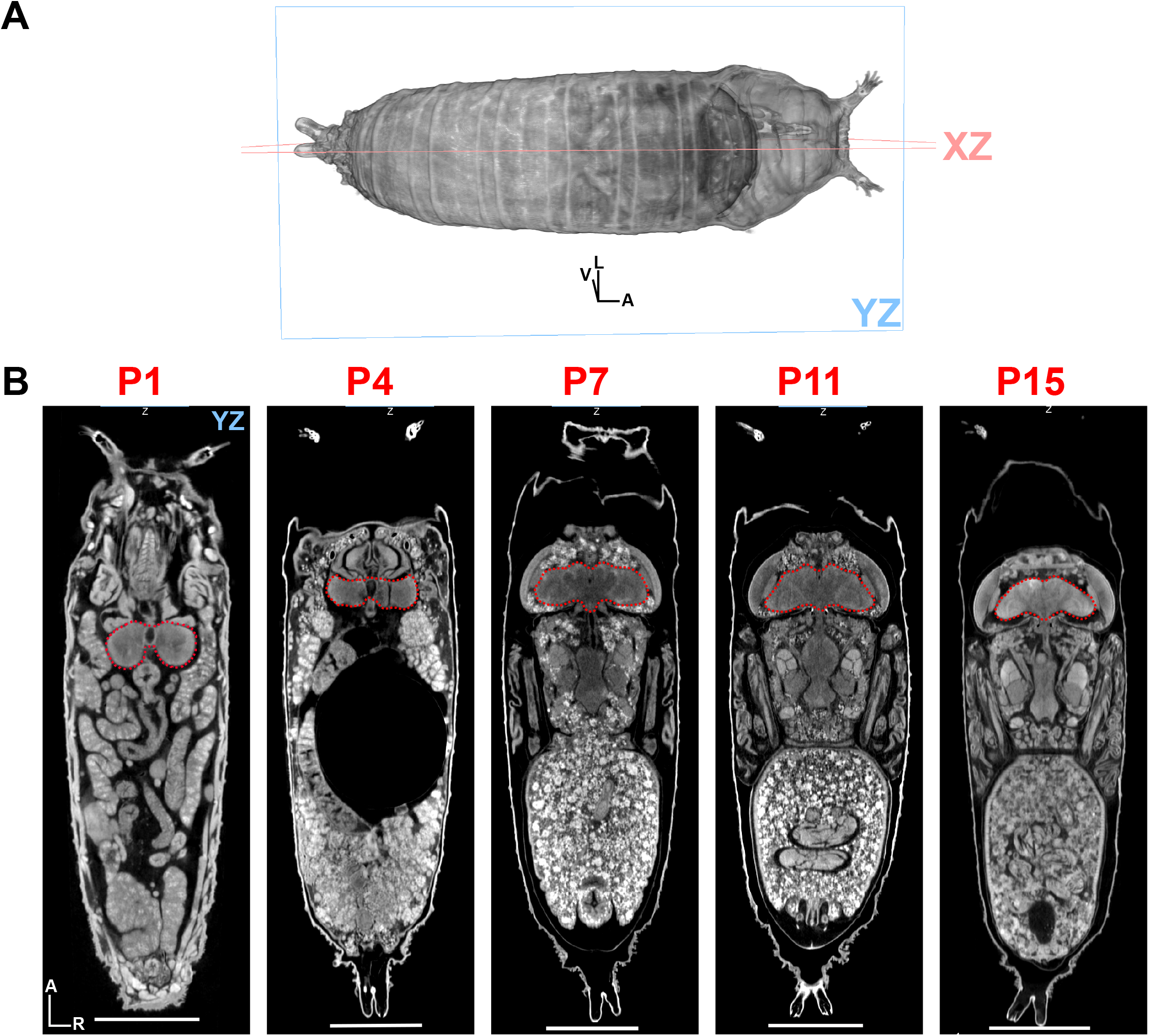
(A) 3D representation of a late stage (P15) pupae, showing body axes (V (Ventral), A (Anterior), L (Left)) and imaging planes XZ and YZ. (B) 2D representation of pupal development at P1, P3/4, P7, P11 and P15 shown in YZ. Brain is outlined in red. Body axes: anterior (A), right (R) Scale Bars (B) = 500 μm.

**Supplementary Figure 3. Related to Figure 7.**
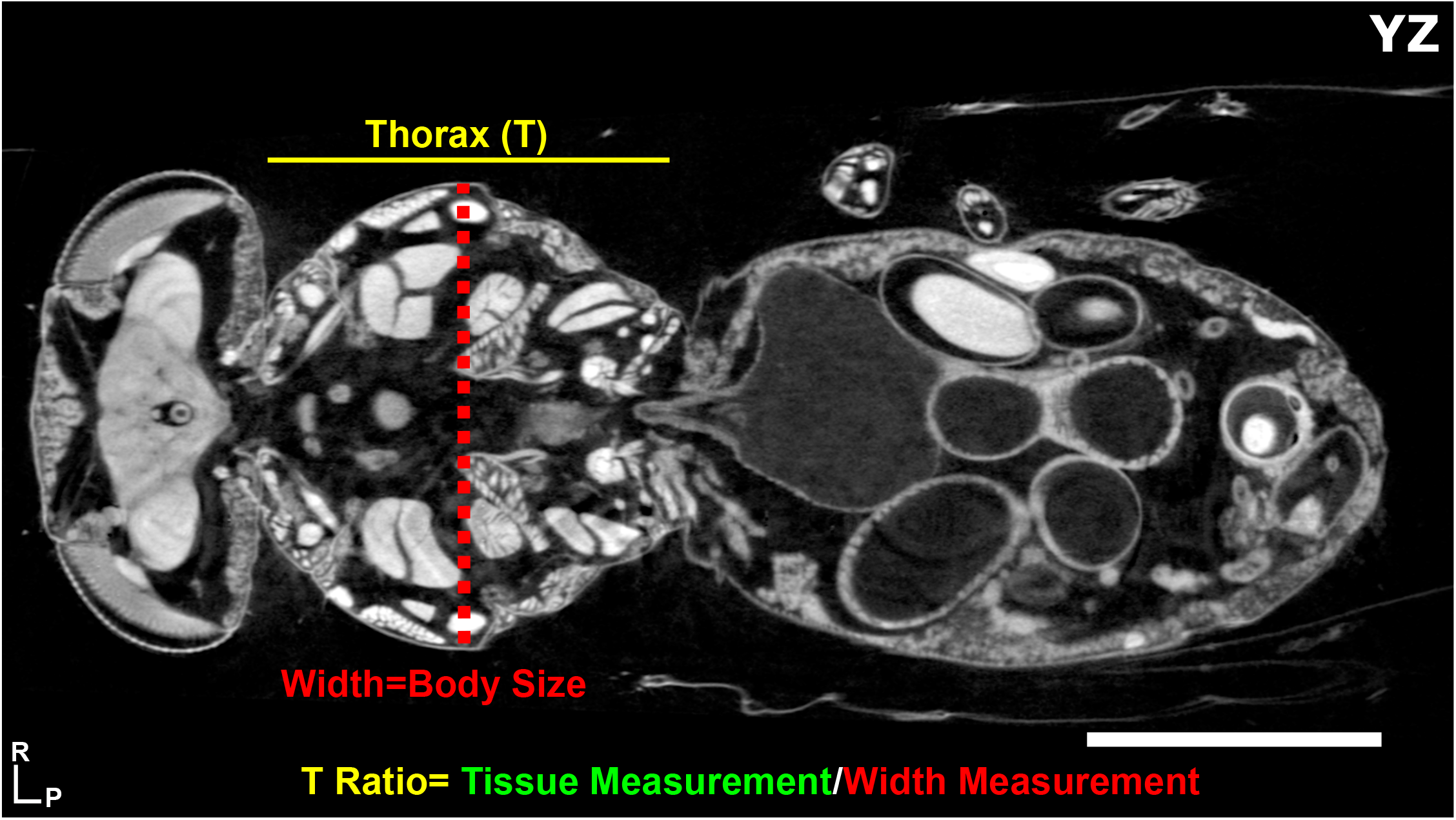
2D YZ view (dorsal perspective) of an adult female fly denoting the thorax width measurement that is used as a proxy for body size normalization to derive the T-Ratio. Body axes: right (R), posterior (P). Scale Bar = 500 μm.

**Supplementary Figure 4. related to Figure 8.**
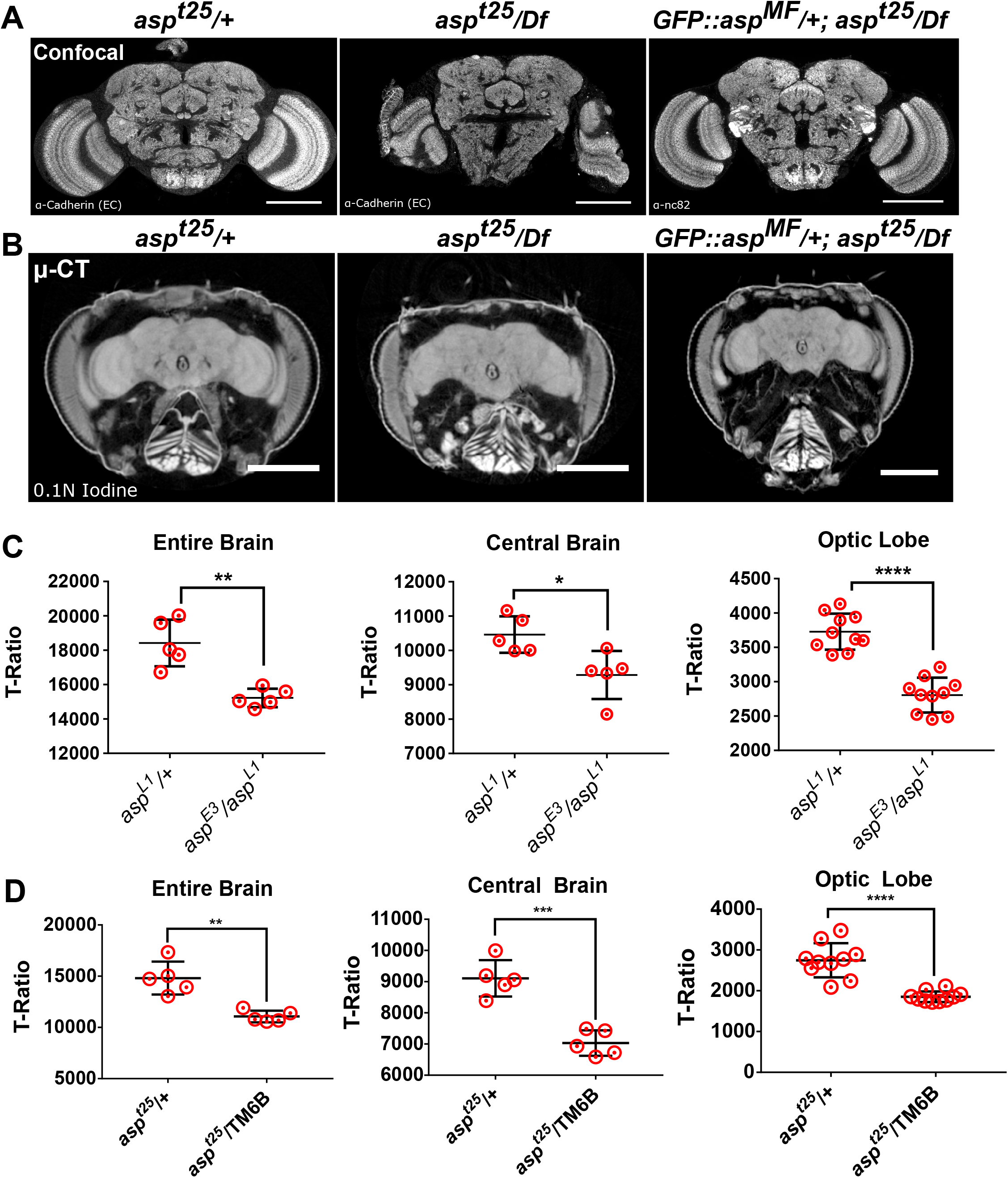
(A) Confocal imaging of control (*asp^t25^*/+), *asp* mutant (*asp^t25^/Df*) and *GFP::asp^MF^* transgenic rescue flies labelled to visualize the neuropil. (B) High resolution (1.2 μm) μ-CT imaging of brains of the same genotype labelled with 0.1N iodine. (C) Volume analysis of the *asp^E3^* and *asp^L1^* alleles, used as a trans-heterozygote. Entire brain, central brain and optic lobe volume expressed as a T-Ratio are shown. (D) Volume analysis of the entire brain, central brain and optic lobes from two control animals, *asp^t25^/+* and *asp^t25^*/TM6B expressed as a T-Ratio. The presence of the balancer chromosome leads to a reduction in brain size, thus, our main analysis in Figure 6 uses *asp^t25^/+* as the control. *n*= 5 brains, unpaired Student’s t-test. *, *P*≤0.05; **, *P*≤0.01; ***, *P*≤0.001; ****, *P*≤0.0001. Scale Bars: (A) 100 μm; (B) 200 μm.

**Supplementary Figure 5. Related to Figure 8.**
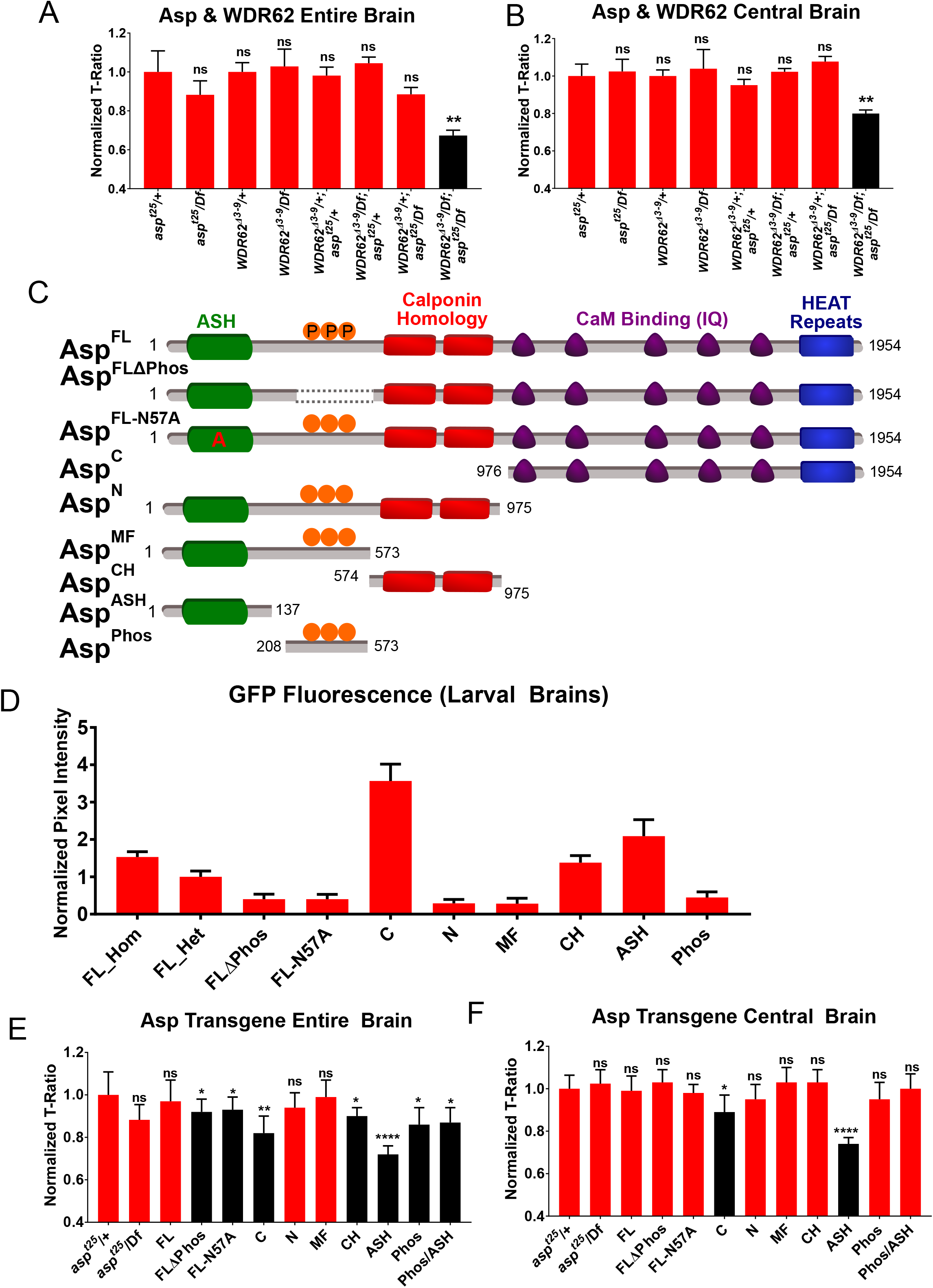
(A) T-Ratio volume analysis of the entire brain from *wdr62 & asp* adult flies. (B) T-Ratio volume analysis of the central brain from *wdr62 & asp* adult flies. (C) Domain representation of the *asp* protein and the various transgenes used in this study. (D) Expression levels of each transgene in the third instar larval brain. Values represent normalized pixel intensity (Methods). (E) T-Ratio volume analysis of the entire brain from *GFP::asp* transgene rescue flies. (F) T-Ratio volume analysis of the central brain from *GFP::asp* transgene rescue flies. *n*= 5 brains, unpaired Student’s t-test. *ns, P*>0.05; *, *P*≤0.05; **, *P*≤0.01; ***, *P*≤0.001; ****, *P*≤0.0001.

**Supplementary Figure 6. Related to Figure 8.**
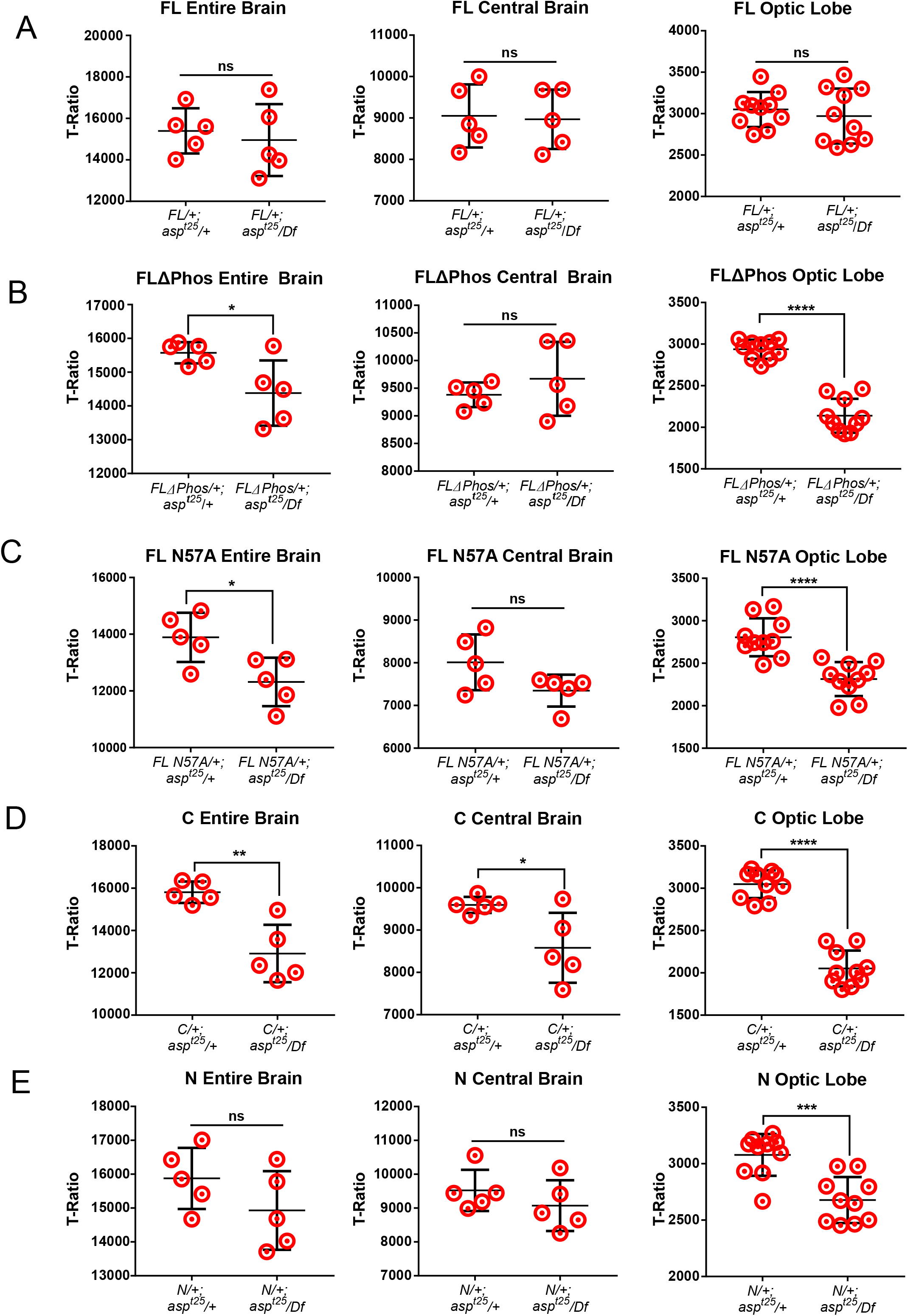
Pairwise comparisons of entire brain, central brain and optic lobe volume expressed as a T-Ratio for each *GFP::asp* transgenic rescue fragment from both control (GFP::asp/+; asp^t25^/+) and mutant (*GFP::asp/+; asp^t25^/Df*) backgrounds. (A) Asp^FL^, (B) Asp^FLΔPhos^, (C) Asp^FLN57A^, (D) Asp^C^, (E) Asp^N^. *n*= 5 brains, unpaired Student’s t-test. *ns, P*>0.05; *, *P*≤0.05; **, *P*≤0.01; ***, *P*≤0.001; ****, *P*≤0.0001.

**Supplementary Figure 7. Related to Figure 8.**
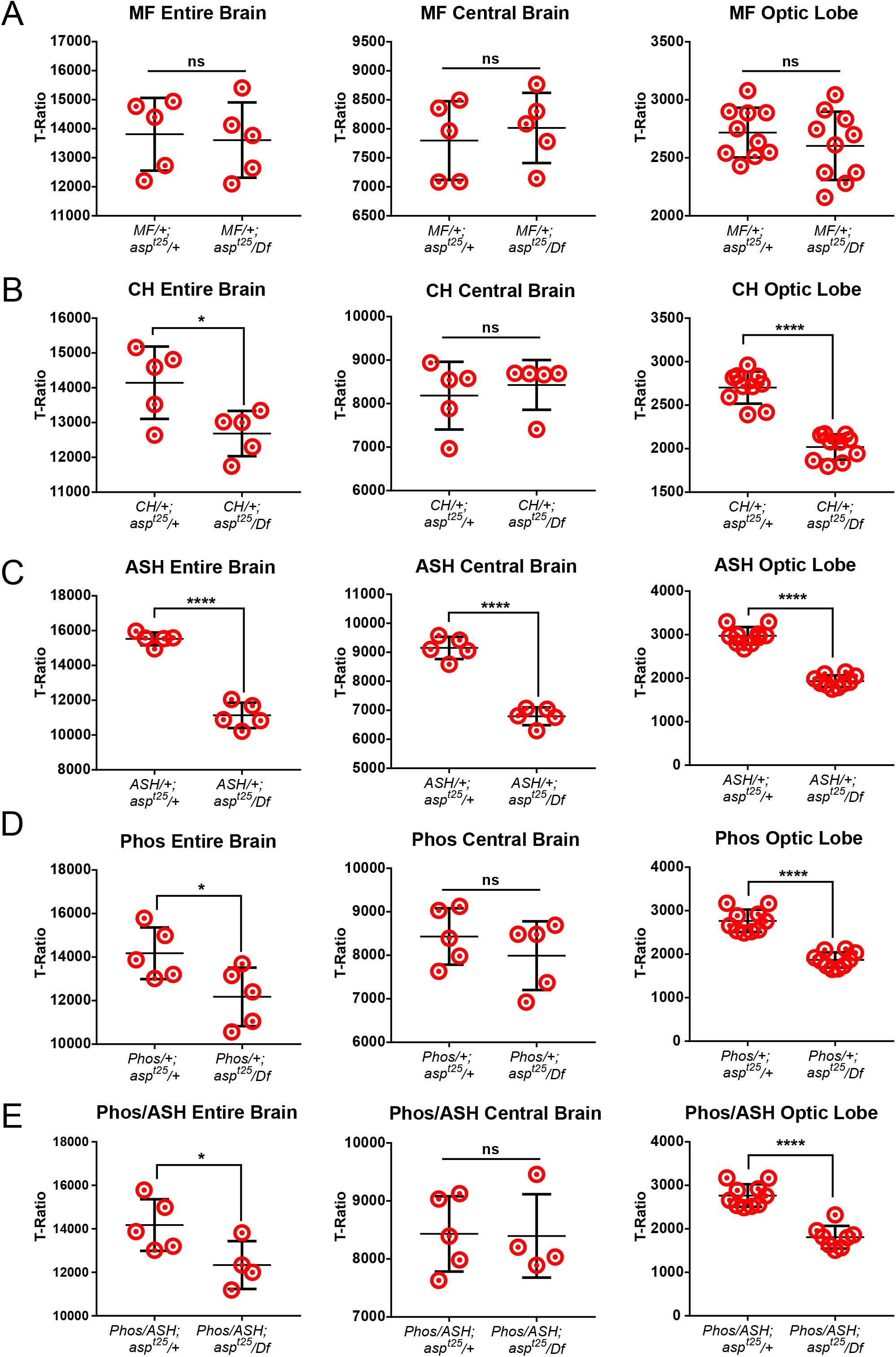
Pairwise comparisons of entire brain, central brain and optic lobe volume expressed as a T-Ratio for each *GFP::asp* transgenic rescue fragment from both control (GFP::asp/+; asp^t25^/+) and mutant (*GFP::asp/+; asp^t25^/Df*) backgrounds. (A) Asp^MF^, (B) Asp^CH^, (C) Asp^ASH^, (D) Asp^Phos^, (E) Asp^Phos/ASH^. *n*= 5 brains, unpaired Student’s t-test. *ns, P*>0.05; *, *P*≤0.05; **, *P*≤0.01; ***, *P*≤0.001; ****, *P*≤0.0001.

## VIDEO LEGENDS

**Video 1. Three-dimensional view of an adult *Drosophila melanogaster* female scanned by μ-CT.** The fly is separated along the XZ (2-14 s) and YZ axes (18-30 s) to reveal internal organs. Individual segmented organs are highlighted as follows: ocelli (red, 39 s); brain (blue, 40 s); lamina (yellow, 41 s); retina (red, 42 s); ventral nerve cord (blue, 43 s); esophagus & foregut (green, 44 s); midgut (yellow, 45 s); hindgut (red, 46 s); boli (purple, 47 s); Malpighian tubules (green, 48 s); crop (lavender; 49 s); salivary glands (turquoise, 50 s); early stage egg chamber (germarium-stage 6, blue, 51 s); mid stage egg chamber (stage 7-11, purple, 52 s); late stage egg chamber (stage 12-14; orange, 53 s); gastrulating embryo (purple, 54 s); heart (blue, 55 s); dorsal longitudinal muscles (orange, 56 s); dorsal ventral muscles (yellow, 57 s; green, 58 s; blue-green, 60 s); jump muscle (orange, 59 s); additional thorax and head muscles (61 s). Stain: 0.1N iodine; 1.25 μm resolution.

**Video 2. Two-dimensional view of an adult *Drosophila melanogaster* female scanned by μ-CT.** The fly is clipped along the X axis to reveal internal structures. Stain: 0.1N iodine; 1.25 μm resolution. Scale Bar: 500 μm.

**Video 3. μ-CT of late third instar larvae.** False-coloring of an intact larvae separated along the XZ axis (0-10 s) and viewed from an anterior-right perspective. 3D rendering of the cuticle and major organ groups: brain (red, 14 s); eye/antennal discs (light blue), leg discs (gold) and haltere discs (yellow) (15 s); wing discs (colored by thickness-see Figure 1, 17 s); gut (foregut, rose; midgut, lavender; hindgut, aqua blue; 18 s); Malpighian tubules (purple, 20 s); fat bodies (blue, 21 s); body wall (green) and mouthpart (yellow) muscles (22 s). Stain: 0.1N iodine; 2 μm resolution.

**Video 4. μ-CT of a wildtype (yw) pupa at stage P11.** The pupa is separated along the YZ axis (5-9 s), followed by clipping along the XY axis from the anterior perspective (15 s). The brain is rendered as a 3D wireframe to reveal its position within the pupal case (18 s). Stain: 0.1N iodine; 1.4 μm resolution.

**Video 5. μ-CT of a wildtype (yw) pupa at stage P15 highlighting the central nervous system.** Position of the ventral nerve cord (VNC, red) and brain (blue) within the thorax and headcase, respectively are highlighted.

**Video 6. RNAi depletion of *spc105r* leads to loss of head structures and pupal lethality, as revealed by μ-CT.** Pupa is initially viewed from the dorsal perspective, anterior is to the right. Position of the ventral nerve cord (VNC, red) and brain (blue) within the thorax is revealed at 10s, the ‘head remnant’ (HR) structure containing an esophagus and labellelum are rendered as a yellow surface. Note orthogonal attachment of brain to VNC within the thorax. Stain: 0.1N iodine; 1.4 μm resolution.

**Video 7. Visualization of the adult thorax muscles by μ-CT.** Adult fly clipped along the XY axis from the anterior perspective to the posterior portion of the thorax. 3D rendering of major muscle thorax muscle groups: dorsal longitudinal muscles (red, 22 s); dorsal ventral muscles (yellow, green, blue-green, 24 s); jump muscle (orange, 24 s); additional thorax muscles (25 s). Stain: 0.1N iodine; 1.25 μm resolution.

**Video 8. Wall thickness of the adult heart.** 3D rendering of the adult heart located dorsally along the abdomen, colored by thickness of the wall (μm) (see Figure 4). Note thickening of the tissue (yellow) at the position of the ostia. Stain: 0.1N iodine; 1.25 μm resolution.

**Video 9. Egg chambers in the adult ovary.** The abdomen is clipped from the dorsal surface to reveal the egg chambers within the ovary. Germarium-stage 6, blue; stage 7-11, yellow; stage 12-14, orange. Stain: 0.1N iodine; 1.25 μm resolution.

**Video 10. An example of left-right asymmetry in adult flies to probe inter-organ relationships.** Directional looping of the male spermiduct (red) over the hindgut (blue). The ejaculatory bulb (green) is shown for reference on the right side of the XZ body axis. Stain: 0.1N iodine; 1.25 μm resolution.

**Video 11. The adult central nervous system.** 3D representation of the ventral nerve cord (VNC, yellow) located along the ventral surface of the thorax and the brain, divided into the central brain (red) and optic lobes (blue). Stain: 0.1N iodine; 1.25 μm resolution.

**Video 12. Two-dimensional view of the adult head imaged at submicron resolution by μ-CT.** XY clip through the head of an adult fly. Stain: 0.5% PTA; ~700 nm resolution. Scale Bar: 100 μm.

**Video 13. The visual circuit from an adult *asp^t25^/+* control fly.** 3D representation of the lamina (green), medulla (blue), lobula (yellow) and lobula plate (red). Stain: 0.5% PTA; 1.2 μm resolution.

**Video 14. The visual circuit from an adult *asp^t25^/Df* mutant fly.** 3D representation of the lamina (green), medulla (blue), lobula (yellow) and lobula plate (red). Note extreme disorganization between the medulla, lobula and lobula plate, severely reduced lamina structures and complete absence of the ocelli. Stain: 0.5% PTA; 1.2 μm resolution.

**Video 15. The visual circuit from an adult GFP::Asp^MF^ transgenic rescue fly.** 3D representation of the lamina (green), medulla (blue), lobula (yellow) and lobula plate (red) from flies expressing Asp^MF^ in the *asp^t25^/Df* mutant background. Note complete rescue of morphology defects and the ocelli. Stain: 0.5% PTA; 1.2 μm resolution.

